# CLAVATA3 peptide signaling from stem cells integrates vascular organization and stem cell maintenance in Arabidopsis shoots

**DOI:** 10.64898/2026.06.25.734304

**Authors:** Krishna Reddy Challa, Caroline A. Sjogren, Nathanaël Prunet, Zachary L. Nimchuk

## Abstract

Terrestrial plant biomass relies on the formation of vascular bundles (VBs) comprising xylem, cambial, and phloem cells that drive secondary growth and wood formation. How VB formation and organization is coordinated with stem growth remains unclear. Here, we show that the conserved CLAVATA3 peptide (CLV3p)-receptor stem cell signaling pathway represses vascular transcriptional programs in the shoot apical meristem (SAM), thereby preventing ectopic vascular differentiation in the stem center, and quantitatively regulates peripheral VB number. Mutational analysis of conserved CLV3p residues partially uncouples its roles in vascular differentiation from stem cell maintenance. However, overexpression and mutational analysis with key phloem regulators reveals that CLV3p controls vascular and stem cell programs via a *DOF* (*DNA-BINDING WITH ONE FINGER*) and *SMXL* (*SUPPRESSOR OF MAX2 1-LIKE*) transcriptional module. Our findings identify a novel role for CLV3p signaling in shoot vascular development and establish a regulatory framework integrating stem cell signaling with vascular patterning.

## Introduction

Understanding how organisms position different tissue types relative to each other during development is an outstanding challenge in biology. Plant vascular tissue forms as strands composed of phloem and xylem tissue, which transport sugar and water respectively and form the stele, along with other associated cell types derived from the procambium.^1–6^ The placement of vascular strands is coordinated with the elaboration of the plant body during development to ensure vascular continuity, water conduction, and solute delivery across plant tissues.^7,8^ In shoots the positioning of vascular strands defines different stelar types and varies across plants species and has informed evolutionary associations in the fossil record.^9–12^ Stele in monocot shoots is dispersed across the stem in a diffuse so-called atactostele pattern. In contrast, shoot stele in dicots like *Arabidopsis thaliana* (Arabidopsis) and gymnosperms are arranged at the stem periphery in a ring, with the center of the stem being composed of pith tissue, which lacks transport functions and is typically parenchymatous.^12–15^ In each dicot vascular strand xylem faces the interior of the stem and phloem faces the exterior, with cambial tissue between. This peripheral stele arrangement, or eustele pattern, is thought to have evolved from an ancestral protostele (haplostele) pattern that is characterized by the presence of a central stele axis and an absence of central pith tissue, as seen in early Devonian fossils from the extinct genera *Psilophyton* and *Cooksonia*.^9,12,14,16^ The mechanisms governing this evolutionary transition are unknown, but the “intrastelar origin” hypothesis posits that pith tissue arose due to the repression of central stele tissue differentiation, therefor restricting stele formation to more peripheral stem regions.^12^

In *Arabidopsis* and other dicots, typically 5–8 vascular bundles form just below the shoot apical meristem (SAM) and later become interconnected through transdifferentiation of interfascicular cambial cells, giving rise to a continuous vascular cambium ring during secondary growth.^1,17–19^ Auxin signaling, together with HD-ZIP III (Class III Homeodomain-Leucine Zipper) transcription factors, specifies xylem identity.^1,20–23^ In parallel, multiple DOF transcription factors and their targets, including SMXL proteins^24^, converge on the master regulator of phloem development APL (ALTERED PHLOEM DEVELOPMENT).^4,25–30^ The peripheral vascular bundles also form in coordination with leaf and floral primordia generation, to ensure new lateral organs are connected to the vascular network.

The consistent production of primordia by the SAM is supported by continual stem cell proliferation at the SAM apex through the CLAVATA3 peptide (CLV3p) signaling pathway.^31–35^ CLV3p signaling is conserved across angiosperms.^36^ CLV3p is produced by the stem cells and diffuses into the central organizing center (OC) cells of the SAM where it is perceived by the CLAVATA1 (CLV1) receptor kinase.^37–39^ CLV1 signaling in the OC dampens the transcription of *WUSCHEL* (*WUS*), which encodes a WOX family homeodomain transcription factor that promotes stem cell proliferation.^40^ WUS in turn promotes *CLV3* expression, forming a feedback loop that maintains stem cell homeostasis.^35,40^ Mutations in *clv1* and *clv3* result in increased SAM stem cell proliferation and meristem size, stem fasciation, and increased floral organ number, all of which can be mimicked by the ectopic expression of *WUS* from the *CLV1* promoter.^41–43^ CLV3p is part of the larger CLE peptide (CLEp) family,^44^ of which there are thirty-three members in Arabidopsis, and CLV1 is part of a subclade of related CLEp receptors which includes BARELY ANY MERISTEM (BAM)1, BAM2 and BAM3.^45,46^ CLV1/BAM receptors require members of the CLE-RESISTANT RECEPTOR KINASE(CLERK)/CLAVATA3 Insensitive Receptor Kinase (CIK) family of co-receptors for signaling.^47–49^ Additional receptors such as the heterodimer of the receptor-like protein CLV2 and the transmembrane pseudokinase CORYNE (CRN) are required for some CLEp functions but are thought to act independently of CLV1/BAM receptors and lack CLEp binding capabilities.^50–53^ Notably, loss of *clv3* across species is associated with transcriptional compensation through upregulation of paralogous *CLEp* genes, which then partially compensate for the loss of *CLV3*.^48,54^ In Arabidopsis, loss of *CLV1* is similarly compensated by transcription of *BAM1-3*, and higher order *clv1;bam1;2;3* quadruple mutants display enhanced stem cell defects compared to *clv1* and *clv3* single mutants.^55^ Transcriptional compensation in *CLV* signaling played a critical role in tomato domestication, by buffering increases in fruit size against extreme stem cell over-proliferation.^54^ Despite this importance to stem cell maintenance and crop domestication, the mechanisms behind *CLV* transcriptional compensation remain unknown.

Other roles for CLEp signaling include promoting fertility, flower primordia formation, and regulating root vascular development.^56–60^ In roots, CLEp signaling through BAM1/2/3 and CLERK/CIKs represses phloem differentiation and the expression of phloem identity genes to spatially organize phloem poles,^58,59,61^ yet BAM1/2 are also required to initiate phloem.^62,63^ In roots, CLEp signaling through BAM3 and CIK2 represses the phloem differentiation and the expression of phloem identity genes.^58,59,61,64^ As neither *bam3* nor phloem *cle* mutants have reported shoot vascular phenotypes, it is unclear if similar or different mechanisms govern shoot vascular identity, formation, and patterning. Here, we show that CLV3p signaling from the SAM stem cells represses phloem-associated gene expression in the SAM center, including the phloem specific paralogue *CLE25*.^48,58^ Correspondingly, *clv3;cle25* double mutant exhibit enhanced SAM defects and also a haplostele-like organization, including ectopic central vascular tissue in the stem. Genetic interaction studies with key vascular regulators establish that CLV3-mediated shoot vascular control depends on the DOF–SMXL transcriptional module. Additionally, ectopic expression of *WUSCHEL* is sufficient to induce vascular differentiation in central pith cells, establishing a CLV3-WUS module in shoot vascular bundle organization. Thus, our work demonstrates that SAM stem cells regulate stele patterning through CLV3 signaling, which simultaneously buffers stem cell proliferation. Furthermore, our results suggest that evolutionary changes in CLV signaling may have contributed to the diversification of stele patterning in flowering plants.

## Results

### CLV3p signaling represses phloem developmental genes in shoot apical meristems

CLV3p signaling maintains stem cell homeostasis through a conserved negative feedback loop that restricts *WUS* expression within the organizing center (OC) (Figure 1A).^65^ While the core architecture of this pathway has been extensively characterized, the downstream transcriptional programs regulated by CLV3p signaling remain poorly understood. To investigate these targets, we established a dexamethasone (DEX)-inducible *CLV3* system in the null *clv3-2* mutant background,^33^ as well as in *clv3-2;clv1-4* plants. *clv1-4* encodes a partially dominant negative receptor variant, which lacks CLEp binding capability.^38,66^ Dominant negative clv1 receptors likely act by interfering with signaling through the related BAM receptors.^67^ As a control, we generated a DEX inducible *mTourquois2* (*mTq2*) line in the *clv3-2* background (Figure 1B). Induction of *CLV3* restored nearly 50% of carpel number toward wild-type levels in *clv3-2*, whereas no rescue was observed in *clv3-2;clv1-4* plants (Figure S1A), confirming that the inducible system is functional, and that CLV3p-dependent regulation of carpel number requires *CLV1* activity.

**Figure 1.**
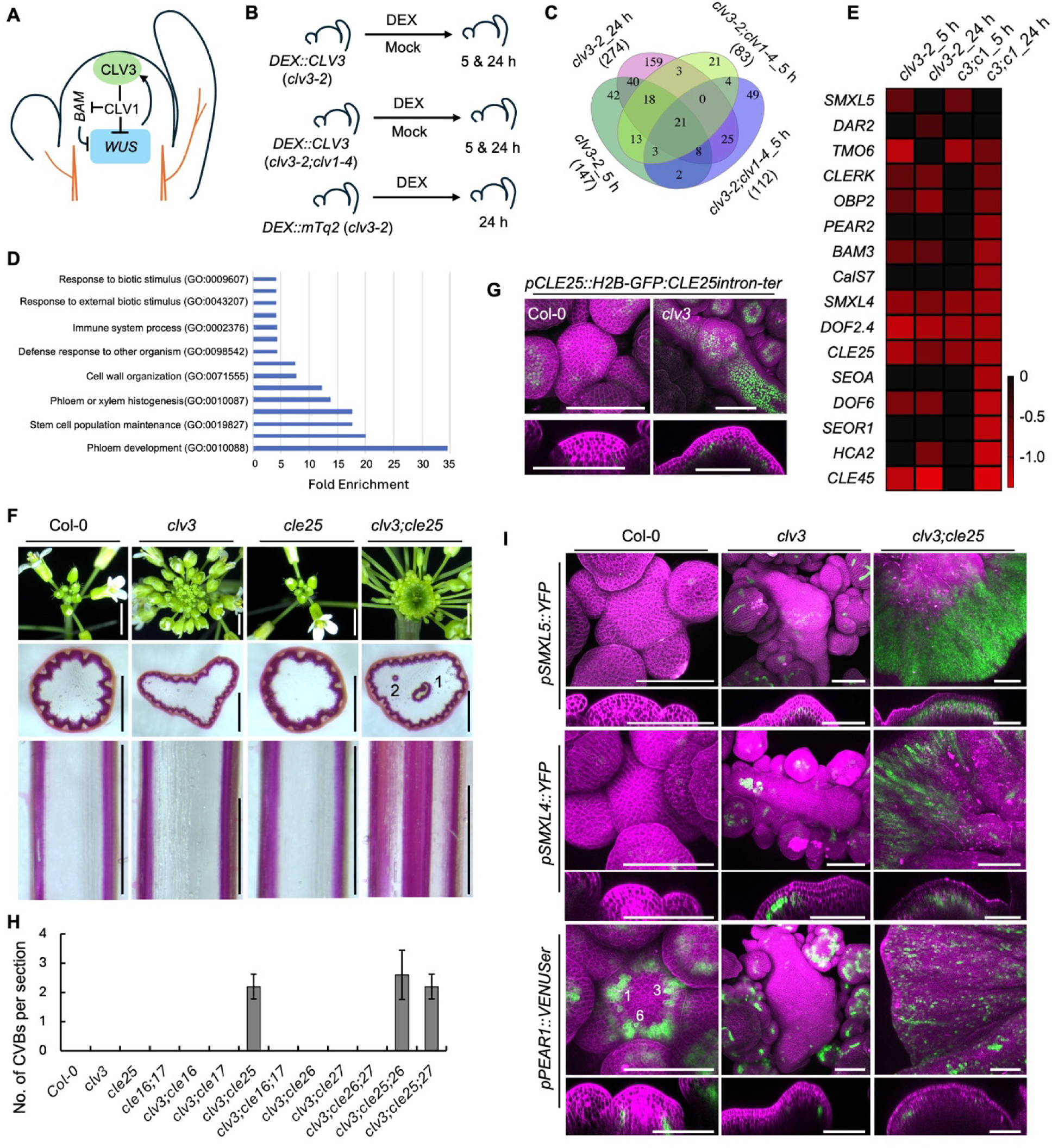
CLV3 and CLE25 peptide signaling redundantly repress phloem-associated gene expression and ectopic vascular formation in the stem center. (A) Schematic representation of the CLV3–WUS regulatory network controlling stem cell maintenance in the shoot apical meristem (SAM). Arrows indicate transcriptional activation, and T-bars indicate transcriptional repression. (B) Experimental design for SAM collection and RNA-seq analysis in *clv3-2* and *clv3-2;clv1-4* backgrounds following dexamethasone (DEX) inducible *CLV3* activation (*DEX::CLV3*) for 5 h or 24 h, together with mock-treated controls. A 24 h DEX-treated sample from *DEX::mTq2*;*clv3-2* plants served as a DEX-system control. *mTq2*; *mTourquois2*. (C) Venn diagram illustrating the overlap among differentially expressed genes identified from the SAM samples described in (B). Numbers indicate differentially expressed gene number. Notably, 21 genes were consistently differentially expressed across all time points. (D and E) Gene Ontology (GO) enrichment analysis of the differentially expressed transcripts shown in (C), highlighting enrichment of phloem-associated genes (D), and corresponding expression profiles of phloem-associated genes displayed as a heatmap (E). The heatmap scale represents log₂ fold-change values. *c3;c1* denotes the *clv3-2;clv1-4* double mutant in (E). (F) Top-view images of SAMs (upper panel), phloroglucinol-stained transverse stem sections (middle), and longitudinal stem sections (bottom) of the indicated genotypes. Numbers indicate CVBs (Central Vascular Bundles) detected in *clv3;cle25* stems. (G) Maximum-intensity projections of the *pCLE25::H2B-int-GFP* reporter in Col-0 and *clv3* SAMs. Representative top-view and longitudinal optical sections are shown. Propidium iodide (PI) staining is shown in magenta. Sample size, n = 3-5. H2B; Histone2B. (H) Quantification of CVB number in Col-0 and the indicated *cle* mutants using the scoring criteria described in (F). Sample size, n = 5. Error bars represent SD. (I) Maximum-intensity projections showing expression of phloem-associated gene transcriptional reporters in SAMs of Col-0, *clv3*, and *clv3;cle25*. Representative top-view and longitudinal optical sections are shown for each reporter line. Propidium iodide (PI) staining is shown in magenta. Sample size, n = 3-5. Scale bars: 2 mm for SAM images in (F), 1 mm for longitudinal sections in (F), 1.5 mm for transverse sections in (F), and 100 μm for panels (G) and (I).

To capture the early transcriptional landscape downstream of CLV3p, we performed a time-course SAM-specific transcriptomic analysis following *CLV3* induction (Figure 1B). In the *clv3-2* background, 147 and 274 genes were differentially expressed at 5 h and 24 h, respectively, with 87 genes shared between the two time points (Figures 1C, S1B and Data S1). In the *clv3-2;clv1-4* background, 83 and 112 genes were differentially expressed at 5 h and 24 h, respectively, with 28 genes shared. Overall, 21 differentially expressed genes were common across both genotypes. The reduced number of responsive genes in the absence of *CLV1* suggests that most CLV3-dependent transcriptional regulation is mediated through CLV1, consistent with the partial inhibition of CLV3p signaling in dominant negative *clv1-4* alleles.

Gene ontology analysis revealed a strong enrichment for phloem developmental regulators among the *CLV3*-repressed genes (Figure 1D). These included multiple members of the *DOF* transcription factor family,^17,29,30,58^ *SMXL* genes,^24,27^ *CLE45*,^63,64^ *CLE25*,^58,68^ and *SIEVE ELEMENT OCCLUSION A* (*SEOA*),^69^ all of which were consistently downregulated at both 5 h and 24 h after *CLV3* induction. Notably, repression of these genes was even stronger in the *clv3-2;clv1-4* background at 24 h (Figure 1E, and Data S1), suggesting that CLV3 signaling suppresses phloem developmental programs through additional receptor pathways, likely involving BAM receptors which overlap with CLV1 in stem cell regulation.^67^ Together, these results uncover an novel role for CLV3 signaling in actively restricting phloem-associated transcriptional identity within the SAM.

### CLV3p functions with CLE25p to repress vascular development in the SAM and stem center

We previously showed that CLV3p functions redundantly with CLE25p in stem cell regulation and floral primordia formation, as *clv3;cle25* double mutants develop enlarged disk-shaped SAMs and exhibit increased carpel number relative to *clv3* single mutants,^48^ whereas *cle25* alone is phenotypically indistinguishable from wild type (Figure 1F). This functional overlap appears to operate largely through transcriptional compensation, with *CLE25* transcription compensating for loss of *CLV3*. Consistent with this, *CLE25* transcripts were rapidly reduced upon *CLV3* induction (Figure 1E), while the *pCLE25::H2B-GFP:CLE25intron-ter* reporter^58^ became ectopically activated throughout the SAM in *clv3* mutants (Figure 1G), indicating that CLV3p suppresses *CLE25* expression in SAMs.

Defects in phloem development are often associated with reduced root and shoot growth.^24,26,70^ Consistent with this, *clv3;cle25* double mutants exhibit a significant reduction in stem height compared with Col-0 (Figure S2A), whereas the *clv3* single mutant shows only a slight decrease. If CLV3p repressed phloem differentiation in SAMs and stems, we suspected that this could be masked by ectopic *CLE25* expression. To investigate this, we sectioned mature first internode stems from Col-0, *clv3*, and *clv3;cle25* genotypes and performed phloroglucinol staining to visualize lignified vascular tissues. Strikingly, *clv3;cle25* stems contain ectopic vascular bundles in the central region, where pith forms (Figure 1F), hereafter termed central vascular bundles (CVBs). On average, *clv3;cle25* stems contained two CVBs per section (Figure 1H). Transverse sections revealed that these CVBs are cylindrical, continuous, and occupy approximately one-quarter of the central stem area. In contrast, CVBs are absent in Col-0, *clv3* and *cle25* plants and vascular in those plants remained restricted to the periphery (Figures 1F and H).

To determine whether CVB formation reflects a general consequence of disrupted CLEp signaling, we analyzed additional *CLE* peptide mutants previously implicated in stem cell maintenance. Although *clv3-15;cle17* and *clv3-15;cle16;17* mutants^71^ developed enlarged disk-like SAMs partially resembling *clv3;cle25*, neither formed CVBs (Figures 1H and S2B). Likewise, mutation of the *CLE25* homologs *CLE26* and *CLE27* failed to induce CVBs in the *clv3* background, and the *clv3;cle25;cle26* triple mutant displayed only a marginal enhancement of the phenotype (Figures 1H and S2C). These observations demonstrate that CLV3p and CLE25p possess a specific and partially redundant function in restricting ectopic vascular bundle formation in stem center. To further elucidate the molecular basis of ectopic CVB formation, we selected the key phloem developmental transcription factors *SMXL4*, *SMXL5*, and *PEAR1* (*PHLOEM EARLY DOF1*), which were strongly repressed upon *CLV3* induction in the RNA-seq dataset (Figure 1E). To assess their spatial expression patterns, we examined promoter reporter activity in the Col-0, *clv3*, and *clv3;cle25* backgrounds. The *pSMXL4::YFP* and *pSMXL5::YFP* signals were undetectable in Col-0 SAMs, whereas both reporters became clearly expressed in the *clv3* mutant (Figure 1I). Owing to the enlarged, disc-shaped SAM morphology of *clv3;cle25*, we defined a confocal imaging region spanning from the central to peripheral zones of the meristem. Both *SMXL4* and *SMXL5* reporters exhibited strong and widespread ectopic expression throughout the SAM in *clv3;cle25* background (Figure 1I).

PEAR genes function as early regulators of vascular bundle development and have been shown to activate SMXL transcription.^7,29,30^ Consistent with previous reports,^7^ *pPEAR1::VENUSer* and *pPEAR2::VENUSer* expression in Col-0 SAMs was preferentially enriched at sites of emerging vascular bundle formation beneath nascent primordia (Figures 1I and S3A). In contrast, in the *clv3* mutant background, *pPEAR1::VENUSer* lost its spatially restricted pattern and instead exhibited widespread punctate expression throughout the SAM. This ectopic expression was further enhanced in the *clv3;cle25* double mutant (Figures 1I and S3A). Notably, this punctate expression pattern was more clearly resolved using the translational reporter *pPEAR2::PEAR2:VENUS*, which accumulated in *clv3;cle25* SAMs but was undetectable in Col-0 and *clv3* backgrounds (Figure S3B), suggesting tight post-transcriptional regulation of PEAR proteins. In addition, another DOF transcription factor reporter, *pOBP2::YFP*, was ectopically expressed in *clv3* SAMs but was not detected in Col-0 (Figure S3C). Collectively, these results indicate that CLV3p, together with CLE25p, suppresses ectopic expression of phloem-associated transcription factors in the SAM center, thereby restricting their expression to discrete domains during vascular bundle formation.

### CLV3 peptide signaling regulates vascular bundle number

*CLV* pathway mutants exhibit progressively stronger acropetal stem cell defects during plant development,^33,54,67^ likely due to cumulative stem cell accumulation. To investigate how this impacts vascular organization, we examined serial stem sections from mature *clv3;cle25;cle27* plants along the basal-to-apical axis (Figure 2A). Phloroglucinol staining revealed a gradual increase in CVB number in an acropetal pattern toward the apex. Basal internodes typically contained only two CVBs, whereas upper internodes closer to the SAM displayed multiple interconnected vascular structures occupying increasingly large portions of the stem center (Figure 2A). These observations reveal that the progressive increase in CVB number mirrors associated with cumulative stem cell accumulation over time.

**Figure 2.**
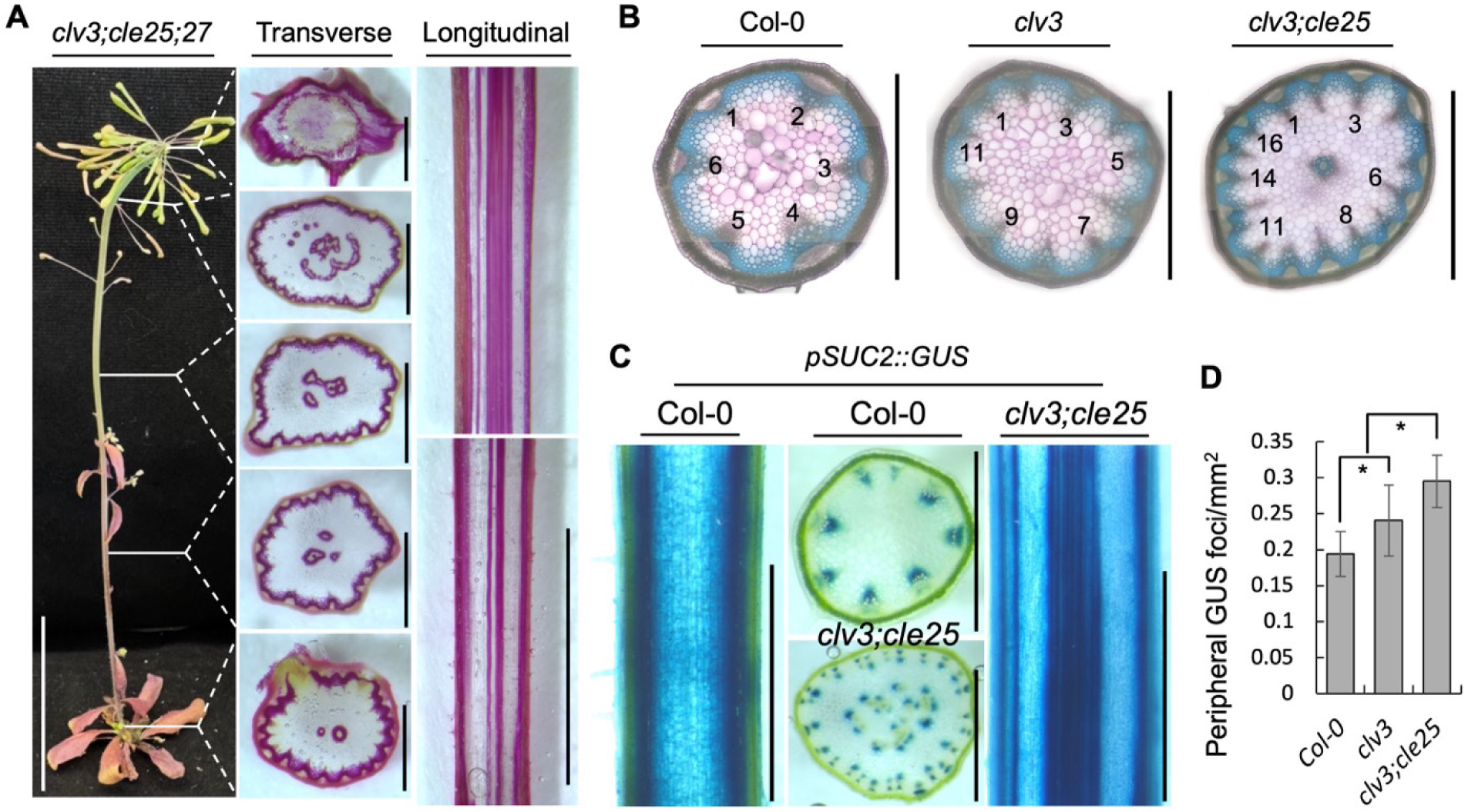
CLV3 peptide signaling maintains vascular bundle number. (A) Two-month-old mature *clv3;cle25;cle27* plants (left), phloroglucinol-stained transverse stem sections collected at regular intervals along the stem axis from basal to apical regions, with sampling positions indicated by white lines (middle), and longitudinal sections of apical and basal stem regions (right). (B) Toluidine blue-stained transverse stem sections of the indicated genotypes. Numbers indicate peripheral vascular bundle counts. Bright blue indicates xylem cells, whereas red-violet marks pectin-rich cells. (C) *pSUC2::GUS* activity in transverse and longitudinal stem sections of the indicated genotypes. (D) Quantification of peripheral GUS-positive foci shown in (C), normalized to stem perimeter. Sample size, n=11-12. Error bars represent SD. Asterisks indicate statistically significant differences (*p* < 0.05), as determined by an unpaired Student’s *t* test. Scale bars: 5 cm for the *clv3;cle25;cle27* plant in (A, left), 1 mm for transverse sections in (A, middle), (B), and (C), 0.5 cm for the longitudinal section in (A, right), and 1.5 mm for longitudinal sections in (C).

Vascular bundles (VBs) are composed of phloem, cambium, and xylem, which together form a characteristic wedge-shaped stele arranged in a circular eustele pattern at the stem periphery in Arabidopsis.^1,15,19^ Toluidine blue staining of Col-0 stem cross-sections clearly visualized these bundles, with typically 5–8 per section (Figure 2B). In contrast, VB number increased to 11 in the *clv3* mutant and further increased to around 16 in *clv3;cle25* stems (Figure 2B). In addition to ectopic CVBs, peripheral vascular bundles in *clv3;cle25* appeared more densely packed than those in Col-0 or *clv3* (Figures 2B and S4A), indicating altered vascular bundle number and spatial patterning in *CLV* pathway mutants.

To quantify these defects, we introduced the *pSUC2::GUS* reporter, which marks phloem companion cells, into the Col-0, *clv3*, and *clv3;cle25* backgrounds (Figure 2C). In Col-0 stems, GUS staining formed a regular peripheral vascular ring with an average density of ∼0.19 foci/mm⁻². Peripheral vascular foci increased significantly in *clv3* (∼0.24 foci/mm⁻²) and further rose to ∼0.29 foci/mm⁻² in *clv3;cle25*, representing nearly a 50% increase over wild type. Strong continuous *pSUC2::GUS* activity was also detected within central CVBs, mimicking the phloroglucinol staining pattern (Figures 1F, 2A and C). Collectively, these findings reveal a novel role for CLV3 peptide signaling in controlling both vascular bundle number and spatial organization in the stem.

### The stem cells of the SAM act as a signaling source for vascular organization

To identify the signaling source that restricts VB number and organization, we expressed *CLV3* or *CLE25* under either the stem cell–specific *CLV3* promoter or the phloem companion cell-specific *SUC2* promoter in the *clv3;cle25* background. Expression of either peptide from the *CLV3* domain fully rescued the enlarged disc-like SAM phenotype and abolished CVB formation (Figure 3A). In addition, stem anatomy and carpel number were restored to near wild-type levels (Figures 3A and B) indicating that the signaling source governing stem cell maintenance is also required to restrict CVB formation.

**Figure 3.**
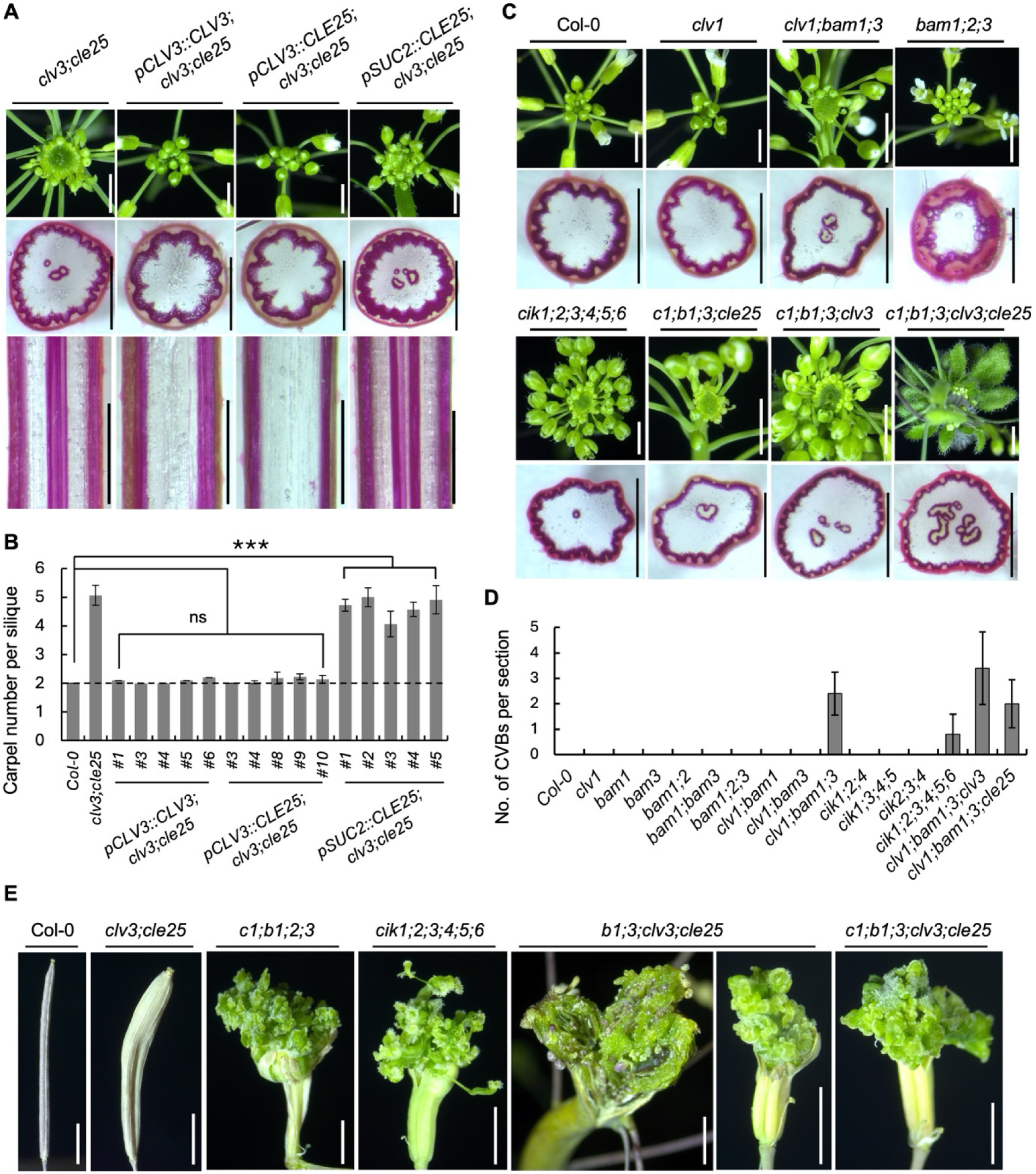
Stem cell derived peptide signaling prevents CVB formation. (A) Top-view images of SAMs (upper panel), phloroglucinol-stained transverse stem sections (middle panel), and longitudinal stem sections (bottom panel) of *clv3;cle25* plants and the indicated transgenic lines. Representative images from five independent T2 lines are shown. (B) Quantification of carpel number per silique in Col-0, *clv3;cle25*, and the indicated transgenic lines. Mean values were calculated from 8–12 siliques collected from 3–5 independent plants per genotype. The dotted line indicates the carpel number in Col-0. Error bars represent SD. Statistical significance was determined using an unpaired Student’s *t* test; \*\*\**p* < 0.001; ns, not significant. (C) Representative SAM images and phloroglucinol-stained transverse stem sections of the indicated CLE receptor mutants and CLE peptide–receptor mutant combinations. (D) Quantification of CVB number in the indicated genotypes. Sample size: n=5 biological replicates per genotype. Error bars represent SD. (E) Representative mature carpels of the indicated genotypes highlighting ectopic proliferation of undifferentiated tissue at carpel apices and stem termini. CVB, central vascular bundle. Scale bars: 2 mm for SAM images in (A, upper panel) and (C), 1 mm for transverse stem sections in (A, middle panel) and (C), 1.5 mm for longitudinal stem sections in (A, bottom panel), and 2.5 mm for (E). *c1, b1, b2*, and *b3* denote *clv1, bam1*, *bam2*, and *bam3*, respectively.

Although CLE25p has been reported to function as a mobile peptide that moves from shoot to root,^72^ phloem-specific expression of *CLE25* under *pSUC2* failed to rescue either the SAM or CVB formation phenotypes. These plants remained indistinguishable from *clv3;cle25* mutants and retained elevated carpel numbers (Figures 3A and B). Together, these results indicate that CLV3 peptide production from the SAM stem cells is essential not only for stem cell regulation, but also for suppression of CVB formation and vascular organization.

### CLE receptors are essential for vascular bundle patterning and organization

CLE peptides regulate stem cell proliferation through multiple receptor complexes, including CLV1 and members of the BAM receptor family. In the absence of *CLV1*, *BAM* receptors become ectopically expressed and partially compensate for stem cell regulatory function.^55,67^ Consistent with this compensatory relationship, CLV3p-mediated repression of phloem-associated genes was enhanced in the *clv1-4* background (Figure 1E), suggesting that CLV3–BAM signaling may play a prominent role in VB organization. To test this possibility, we generated a series of higher-order *CLE* receptor mutant combinations and examined their vascular phenotypes. Neither single nor double mutant combinations of *CLV1* and *BAM1-3* receptors developed ectopic CVBs formation (Figures 3C, D, and S4B), and stem morphology remained largely comparable to that of Col-0. In contrast, the *clv1;bam1;bam3* triple mutant developed prominent CVBs together with a disk-like SAM phenotype closely resembling *clv3;cle25* (Figures 3C and D). Interestingly, although *bam1;bam2;bam3* triple mutants displayed altered stem morphology, including increased stem thickness, they failed to develop CVBs.

A similarly high degree of functional redundancy was observed among the CLERK/CIKs, which are CLV1/BAM co-receptors.^47,49,58,59,73^ Whereas lower-order *cik* mutant combinations, including the triple mutants (*cik1;2;4* and *cik2;3;4*) and the quadruple mutant (*cik1;3;4;5*), showed no detectable vascular defects, the *cik* sextuple mutant (*cik1;2;3;4;5;6*) developed low-frequency CVBs accompanied by a disk-like SAM phenotype (Figures 3C, D, and S4B). Collectively, these observations reveal extensive functional redundancy among CLE peptide receptors and co-receptors in suppressing ectopic CVB development.

The CVB phenotype became substantially more severe when receptor mutations were combined with *clv3* and *cle25* peptide mutants (Figures 3C, D, and S4B). Introduction of *clv3*, but not *cle25*, into the *clv1;bam1;bam3* background dramatically enhanced CVB formation, indicating that CLV3 is the predominant ligand controlling VB patterning (Figures 3C and D). The higher-order *clv1;bam1;3;clv3;cle25* mutant developed enlarged and extensively interconnected CVBs throughout the stem (Figure 3C), precluding reliable quantification of individual CVB numbers. Consistent with these phenotypic abnormalities, RT–qPCR analysis revealed strong upregulation of phloem-associated developmental regulators, including *SMXL5*, *PEAR2*, and *HCA2* (*HIGH CAMBIAL ACTIVITY 2*), in higher order receptor mutant combinations (Figure S4C).

Beyond their vascular abnormalities, these peptide–receptor mutant combinations exhibited severe stem splitting and massive proliferation of undifferentiated tissue at stem and floral apices during late stages of development, phenotypes absent in Col-0 and very rarely observed in *clv3;cle25* plants (Figures 3E and S5). These defects are consistent with severe deregulation of WUS-dependent stem cell activity and were strongly associated with the extent of ectopic CVB formation. Together, these results reveal that CLE peptide–receptor signaling complexes, beyond their established roles in stem cell maintenance, are also essential for proper vascular bundle patterning and spatial organization in stems.

### Decoupling the dual functions of CLV3 peptide signaling

Our *CLV3* induction transcriptome revealed that CLV3p strongly represses *CLE45* expression (Figure 1E). In Arabidopsis roots, the CLE peptides CLE33 and CLE45 redundantly inhibit phloem differentiation through an autocrine signaling pathway mediated primarily by the BAM3 receptor.^74,75^ To determine whether these peptides functionally interact with CLV3 signaling during shoot vascular development, we generated higher-order peptide mutant combinations, including the *clv3;cle33;45* triple mutant and the *clv3;cle25;33;45* quadruple mutant. The triple mutant largely phenocopied *clv3* (Figures 1F and 4B), the quadruple mutant exhibited an enlarged, disk-like SAM together with mild increased CVB formation compared with *clv3;cle25* (Figure 4A). These observations suggest that other phloem-associated CLE peptides contribute modestly to CLV3p-dependent vascular organization.

**Figure 4.**
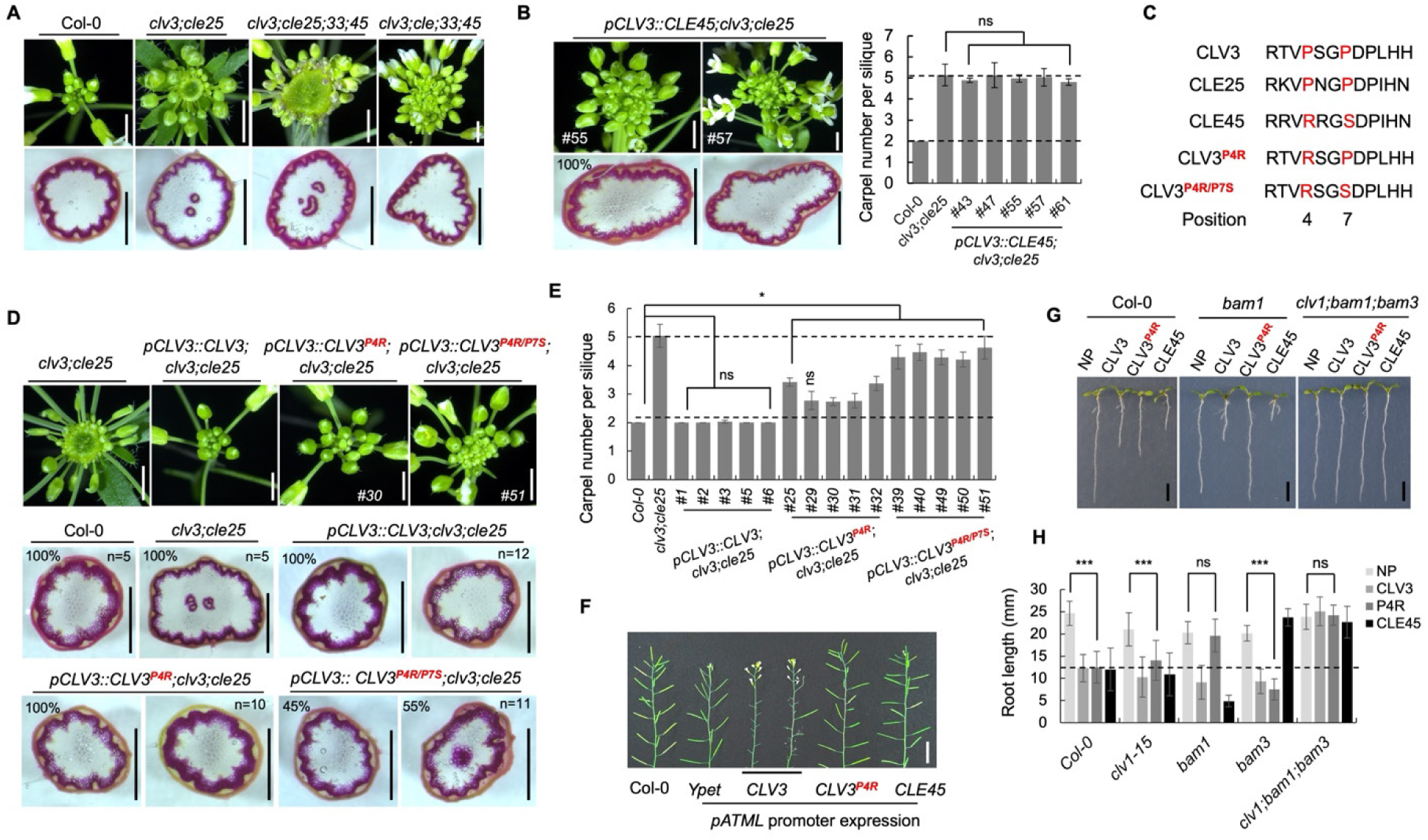
Engineered CLV3 peptide variants uncouple stem cell maintenance from suppression of CVB formation. (A) Top-view images of SAMs (upper panel) and phloroglucinol-stained transverse stem sections (bottom panel) of the indicated genotypes. (B) Representative SAM images (upper panel) and phloroglucinol-stained transverse stem sections (bottom panel) of two independent *pCLV3::CLE45;clv3;cle25* transgenic lines (#55 and #57; left). Quantification of carpel number per silique in Col-0, *clv3;cle25*, and five independent *pCLV3::CLE45;clv3;cle25* lines is shown at right. Error bars represent SD. Mean values were calculated using 8–12 siliques from 3–5 independent plants per genotype. For CVB analysis, 13 independent T1 lines and 5 T2 lines were examined, and representative images are shown. (C) Sequence alignment of CLE domains from the indicated CLE peptides. Conserved residues at positions 4 and 7 are highlighted in red. Modified CLV3 peptide variants carrying substitutions at position 4 (CLV3^P4R^) or at positions 4 and 7 (CLV3^P4R/P7S^) are indicated. (D) Representative SAM images (top panel) and phloroglucinol-stained transverse stem sections (bottom panel) of the indicated transgenic lines expressing wild-type or modified CLV3 peptide variants shown in (C), together with *clv3;cle25* controls. Percentages of plants exhibiting CVB formation and the corresponding numbers of independent T1 lines (n) are indicated. (E) Average carpel number per silique in the transgenic lines shown in (D), compared with Col-0. Mean values were calculated using 8–12 siliques from 3–5 independent T2 plants per genotype. Error bars represent SD. (F) Inflorescences showing silique formation in transgenic plants expressing *CLV3*, *CLV3^P4R^, or CLE45* under the *pATML* promoter in the Col-0 background, together with *Ypet* vector controls. (G and H) 7-day-old seedlings (G) and quantification of primary root length (H) of the indicated genotypes grown on one-half-strength MS medium without peptide (NP) or supplemented with 0.1 μM CLV3, CLV3^P4R^, or CLE45 synthetic peptides. Quantification in (H) compares root lengths of NP- and CLV3^P4R^-treated (P4R) seedlings. Sample size, n = 13–15. Error bars represent SD. Statistical significance was determined using an unpaired Student’s *t* test; \**p* < 0.05, \*\*\**p* < 0.001; ns, not significant. The bottom and top dotted lines in (B) and (E) indicate the carpel numbers in Col-0 and *clv3;cle25*, respectively. Scale bars: 2 mm for SAM images in (A), (B), and (D); 1 mm for transverse sections in (A), (B), and (D); 3 mm for (F); and 5 mm for (G).

To directly test the function of CLE45p in stem vascular organization, we ectopically expressed *CLE45* under the *CLV3* promoter in the *clv3;cle25* background. Interestingly, *pCLV3::CLE45* strongly suppressed ectopic CVB formation and partially restored SAM morphology. However, carpel number remained largely unchanged (Figures 4B and S6), and these transgenic plants closely resembled *clv3* mutants (Figures 1F and 4B), suggesting that CLE45p preferentially regulates vascular organization while contributing only weakly to stem cell homeostasis.

To understand the molecular basis underlying this functional divergence, we compared the CLE peptide domains of CLV3, CLE25, and CLE45 (Figure 4C). Sequence analysis identified two amino acid substitutions at highly conserved positions within the CLE motif. Whereas CLV3p and CLE25p contain proline (P) residues at positions 4 and 7, CLE45p instead carries arginine (R) and serine (S) residues at these positions, respectively (Figure 4C). We hypothesized that these substitutions may selectively alter downstream signaling outputs, possibly by discriminating among CLV1/BAM receptors.^63,76,77^ To test this idea, we engineered modified CLV3 peptides carrying either a single P4R substitution (*CLV3^P4R^*) or combined P4R/P7S substitutions (*CLV3^P4R/P7S^*) and expressed them under the native *CLV3* promoter in the *clv3;cle25* background.

Strikingly, the single P4R substitution was sufficient to fully suppress CVB formation, comparable to wild-type *CLV3* transgenes. However, these plants retained moderately enlarged SAMs and elevated carpel numbers (Figures 4D and E), indicating they retain partial loss of stem cell regulatory activity. The double-substitution variant (*CLV3^P4R/P7S^*) failed to restore carpel number to wild-type levels, despite still suppressing ectopic CVB formation in more than 50% of transgenic lines (Figures 4D and E). These observations suggest that vascular bundle organization and stem cell maintenance can be genetically decoupled in part through discrete modifications within the CLE peptide sequence. Consistent with this interpretation, ectopic expression of wild-type *CLV3* under the epidermal specific *ATML1* promoter^78^ is sufficient to terminate carpel development in Col-0 plants, as previously reported.^79^ By contrast, expression of *CLV3^P4R^*or *CLE45* under epidermal specific *ATML1* promoter failed to induce floral termination and instead produced plants indistinguishable from vector controls (Figure 4F). Together, these results support a model in which discrete residues within the CLE domain differentially modulate downstream signaling outputs, leading to a partial decoupling of CLV3-mediated regulation of vascular bundle formation and stem cell homeostasis.

### CLV3^P4R^ peptide function through BAM1

Previous studies have demonstrated that synthetic CLV3 and CLE45 peptides inhibit primary root growth by promoting premature differentiation of root meristematic cells.^64^ To determine the receptor specificity underlying CLV3^P4R^-mediated signaling, we synthesized CLV3, CLE45, and CLV3^P4R^ peptides and examined their effects on primary root growth in different receptor mutant backgrounds (Figures 4G and H). Consistent with earlier reports, CLV3p-mediated root growth inhibition was not much altered in the single mutants *clv1*, *bam1*, and *bam3*, but was almost completely abolished in the *clv1;bam1;3* triple mutant,^63,76,80^ indicating that CLV3p signals redundantly through multiple receptors in roots. In contrast, CLE45p-mediated root growth inhibition was specifically compromised in the *bam3* mutant (Figure 4H), confirming the preferential requirement of BAM3 for CLE45p signaling.^64^

However, CLV3^P4R^-mediated root growth inhibition was selectively attenuated in the *bam1* mutant background (Figures 4G and H), while remaining largely unaffected in *clv1* or *bam3*. These results indicate that the P4R substitution redirects CLV3p signaling toward a BAM1-dependent pathway. This receptor preference is consistent with the enhanced repression of phloem-associated genes following *CLV3* induction in the *clv1-4* background (Figure 1E), where BAM receptors are ectopically expressed.^55,67^ Together, these data suggest that subtle changes within the CLE peptide sequence can reprogram receptor utilization and thereby alter developmental output specificity.

### Phloem DOF transcription factors act downstream of CLV3 signaling to coordinate stem cell homeostasis and vascular bundle patterning

Phloem-associated DOF transcription factors function within a highly redundant regulatory network. Among them, *PEAR1* (*DOF2.4*) and *PEAR2* (*DOF5.1*) are strongly enriched in developing protophloem (Figure 5A), while additional family members including *DOF3.2*, *DOF5.3*, *OBP2* (*DOF1.1*), and *HCA2* (*DOF5.6*) act cooperatively to promote vascular development.^25,29,58^ Consistent with this redundancy, single loss-of-function mutants exhibit little or no visible phenotype, whereas higher-order *dof*-sextuple mutants (*obp2;dof2.2;pear1;pear2;dof5.3;dof3.2*)^58^ develop severe small-root phenotypes. Our transcriptomic and reporter analyses revealed that multiple DOF genes, including *PEAR1*, *PEAR2*, *OBP2*, and *HCA2*, are coordinately repressed together with *SMXL* genes following *CLV3* induction (Figures 1C-E, I and S3), suggesting that PEARs function as early downstream targets of CLV3p signaling during vascular organization. To test this hypothesis, we introduced CRISPR/Cas9-generated *clv3* and *cle25* mutations into the published *dof*-sextuple^58^ background using the same guide RNAs previously our lab employed to generate the *clv3;cle25* line^48^. *Cas9*-free *dof-sextuple;clv3* and *dof-sextuple;clv3;cle25* lines carrying the corresponding edited *CLV3* and *CLE25* alleles were then verified by sequencing.

**Figure 5.**
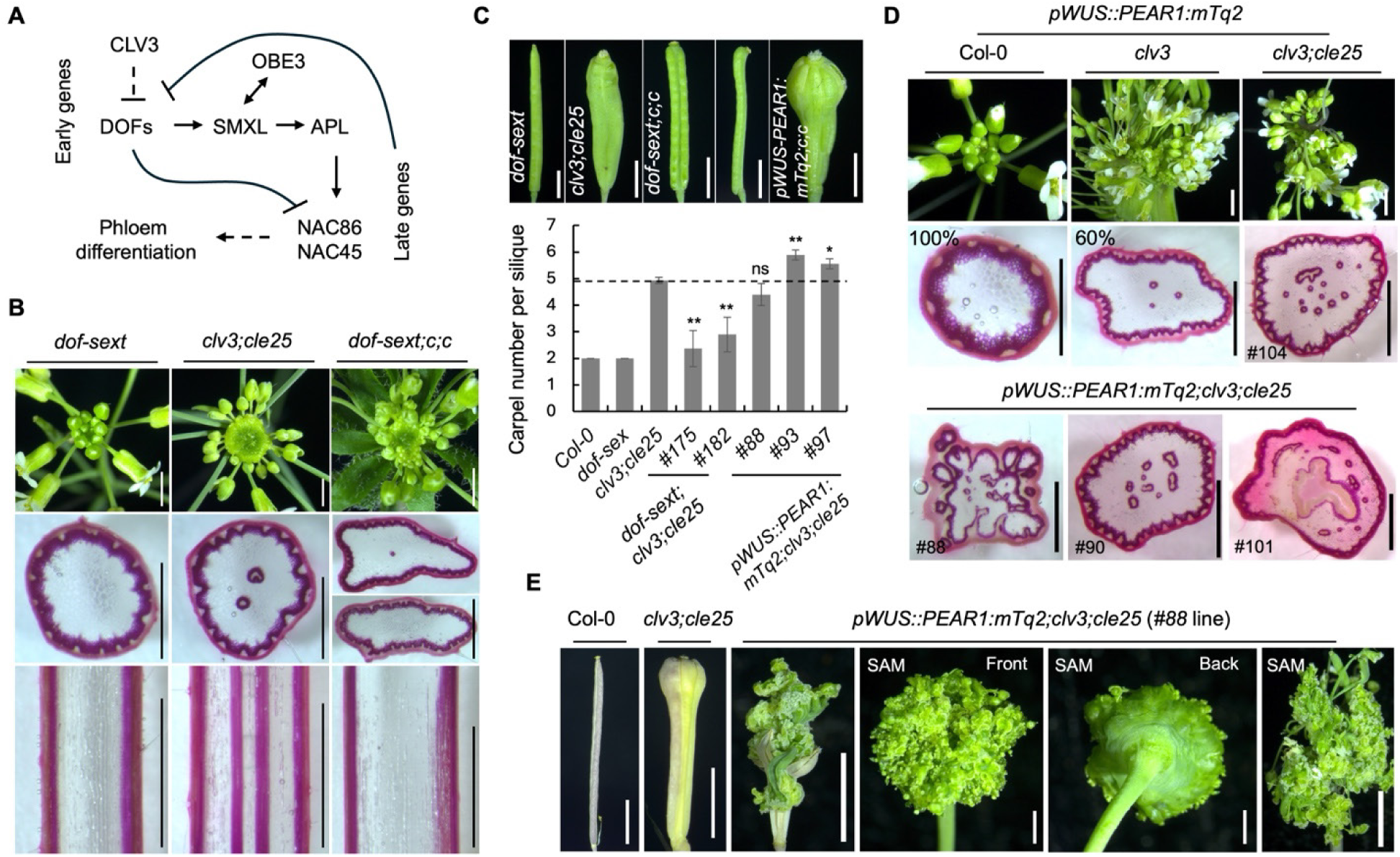
Phloem-associated DOF transcription factors integrate CLV3 signaling in stem cell maintenance and CVB suppression. (A) Schematic model of the established phloem-associated DOF transcriptional network regulating phloem differentiation. Arrows indicate transcriptional activation, T-bars indicate transcriptional repression. Dashed arrows denote indirect regulatory interactions through intermediate factors. (B) SAM images (top panel), phloroglucinol-stained transverse stem sections (middle panel), and longitudinal stem sections (bottom panel) of *dof*-sextuple (*dof*-sext), *clv3;cle25*, and *dof*-sext;*clv3;cle25* (*dof*-sext;*c;c*) plants. A small fraction of *dof-sext;clv3;cle25* stem cross-sections exhibited a single dot-like vascular bundle focus in the stem center. (C) Silique images (top panel) and quantification of the average carpel number per silique (bottom panel) in the indicated genotypes. *c;c* denotes the *clv3;cle25* genotype in the silique images. Quantification was performed using 8–12 siliques from 3–5 independent plants per genotype. The dotted line indicates the average carpel number of *clv3;cle25*. Error bars represent SD. Statistical significance was determined using an unpaired Student’s t-test. Statistical analyses were performed relative to *clv3;cle25*. *P < 0.05, **P < 0.005, and ns indicates not significant. (D) Top-view SAM images (top panel) and phloroglucinol-stained transverse stem sections (bottom panels) of the indicated genotypes. Percentages of plants with and without central vascular bundle (CVB) formation are indicated. Sample size, n=5–12. Independent transgenic line numbers are shown. (E) Representative mature siliques (left) and floral SAM termini (right) of the indicated genotypes, highlighting ectopic proliferation of undifferentiated tissue in *pWUS::PEAR1:mTq2* expressing lines Scale bars: 2 mm for SAM images in (B), and (D), and siliques in (C) and (E); 1 mm for transverse stem sections in (B) and (D); and 2 mm for longitudinal stem sections in (B).

The stems of *dof-sextuple;clv3* mutants were frequently widened and occasionally bifurcated, with no CVB formation (Figure S7A and B). The octuple mutant *dof-sextuple;clv3;cle25* retained only a weak disk-like SAM phenotype, and CVB development was almost entirely abolished, with only rare phloem-associated puncta detectable in a subset of stems (Figure 5B). Histochemical analyses of longitudinal stem sections further confirmed the near-complete suppression of ectopic vascularization (Figure 5B). These findings demonstrate that CLV3p-mediated vascular patterning is dependent on phloem-associated DOF transcription factors.

Surprisingly, both the *dof-sextuple;clv3* and *dof-sextuple;clv3;cle25* mutants exhibited a strong reduction in carpel number relative to the *clv3* and *clv3;cle25* controls (Figures 5C and S7C). Consistent with our previous observations,^48^ *clv3;cle25* plants produced an average of 4.9 carpels per silique, with approximately 30% of siliques containing five or six carpels. In contrast, *dof-sextuple;clv3;cle25* plants produced an average of 2.5-3 carpels per silique, with 47% of siliques displaying the two carpels and around 10% containing four carpels (Figure 5C). A similar restoration of carpel number was observed in *dof-sextuple;clv3* plants (Figure S7C), further supporting a critical role for DOF transcription factors in maintaining stem cell homeostasis downstream of CLV3p signaling.

### PEAR1 promotes stem cell proliferation

To determine whether ectopic DOF activity is sufficient to disrupt stem cell maintenance in shoots, we expressed *mTq2*-tagged *PEAR1* under the control of the *WUS* promoter (*pWUS::PEAR1:mTq2*) in Col-0, *clv3*, and *clv3;cle25* backgrounds (Figure 5D). *PEAR1* overexpression in Col-0 resulted in marginally enlarged SAM in few plants, but the majority were indistinguishable from wild type. In contrast, *PEAR1* strongly enhanced stem cell–associated phenotypes in both *clv3* and *clv3;cle25* mutants, promoting meristem fasciation and inward curling of the SAM (Figure 5D). In the *clv3;cle25* background, carpels and rosette branches accumulated excessive undifferentiated tissue at their apices, giving rise to enlarged disk-like structures (Figure 5E). These abnormalities closely resembled the phenotypes of higher-order CLE peptide–receptor mutants (Figures 3E and S5), in which phloem *DOF* and *SMXL* genes are strongly upregulated (Figure S3C), thereby linking phloem-associated transcriptional programs to the regulation of stem cell homeostasis.

Consistent with the enhanced meristem defects, *PEAR1* overexpression in the *clv3;cle25* background further enhanced the carpel number phenotype in two of three independent transgenic lines (Figure 5C). Approximately 41% of siliques developed five carpels, whereas 30% produced seven carpels, with rare siliques containing up to nine carpels, a phenotype not observed in *clv3;cle25* plants. In contrast, ectopic expression of another phloem-associated DOF transcription factor, *DOF2.2*, under the *STM* promoter failed to induce detectable phenotypic changes in either *clv3* or *clv3;cle25* backgrounds (Figure S8), indicating a specific role for PEAR1 in promoting stem cell proliferation. Together, these results revealed that phloem-associated DOF transcription factors are key effectors of CLV3p signaling in the regulation of stem cell homeostasis.

### PEARs are essential for vascular organization in CLV3 signaling

In addition to regulating stem cell homeostasis, ectopic expression of *PEAR1* within the *WUS* domain (*pWUS::PEAR1:mTq2*) was sufficient to induce ectopic CVB formation in three out of five independent transgenic lines in the *clv3* background (Figure 5D), where control *clv3* plants never developed CVB (Figure 1F and H). This phenotype became substantially more severe in *clv3;cle25*, with stems frequently producing 8–10 ectopic CVBs. Moreover, strong *pWUS::PEAR1:mTq2;clv3;cle25* lines displayed dramatic disorganization of peripheral vascular bundle architecture in addition to extensive ectopic central vascularization (Figure 5D), indicating that strict spatial repression of DOF activity within the organizing center is essential for maintaining normal vascular patterning.

### *SMXLs* act downstream of CLV3 signaling to control SAM growth and vascular patterning

Our transcriptomic and reporter analyses revealed that CLV3 signaling strongly represses *SMXL* gene expression in the SAM (Figures 1E and I), whereas SMXL proteins are established positive regulators of phloem development that function downstream of DOF transcription factors (Figure 5A).^24,27–30^ Based on these observations, we hypothesized that CLV3p-dependent vascular bundle patterning is mediated through suppression of *SMXL* expression.

To test this, we introduced the *smxl4;smxl5* double mutant into the *clv3;cle25* background. Consistent with previous reports,^24,81^ *smxl4;smxl5* plants exhibited severe root growth defects associated with impaired phloem formation, while the SAM remained largely unaffected (Figure 6A). In contrast, *clv3;cle25;smxl4;smxl5* quadruple mutants displayed synergistic growth defects, including severe developmental retardation and excessive anthocyanin accumulation in rosette leaves (Figure S9A). Despite these defects, the quadruple mutants retained the characteristic disk-shaped SAM observed in *clv3;cle25* plants, although meristem size was markedly reduced (Figures 6A-C). Quantitative measurements revealed that SAM area decreased from around 2 mm² in *clv3;cle25* plants to ∼0.5 mm² in *clv3;cle25;smxl4;smxl5* mutants. Similarly, floral bud number was reduced in both *clv3;smxl4;smxl5* and *clv3;cle25;smxl4;smxl5* backgrounds, whereas carpel number remained largely unaffected (Figures 6A and S9B). Collectively, these observations indicate that SMXLs contribute to stem cell proliferation downstream of CLV3 signaling.

**Figure 6.**
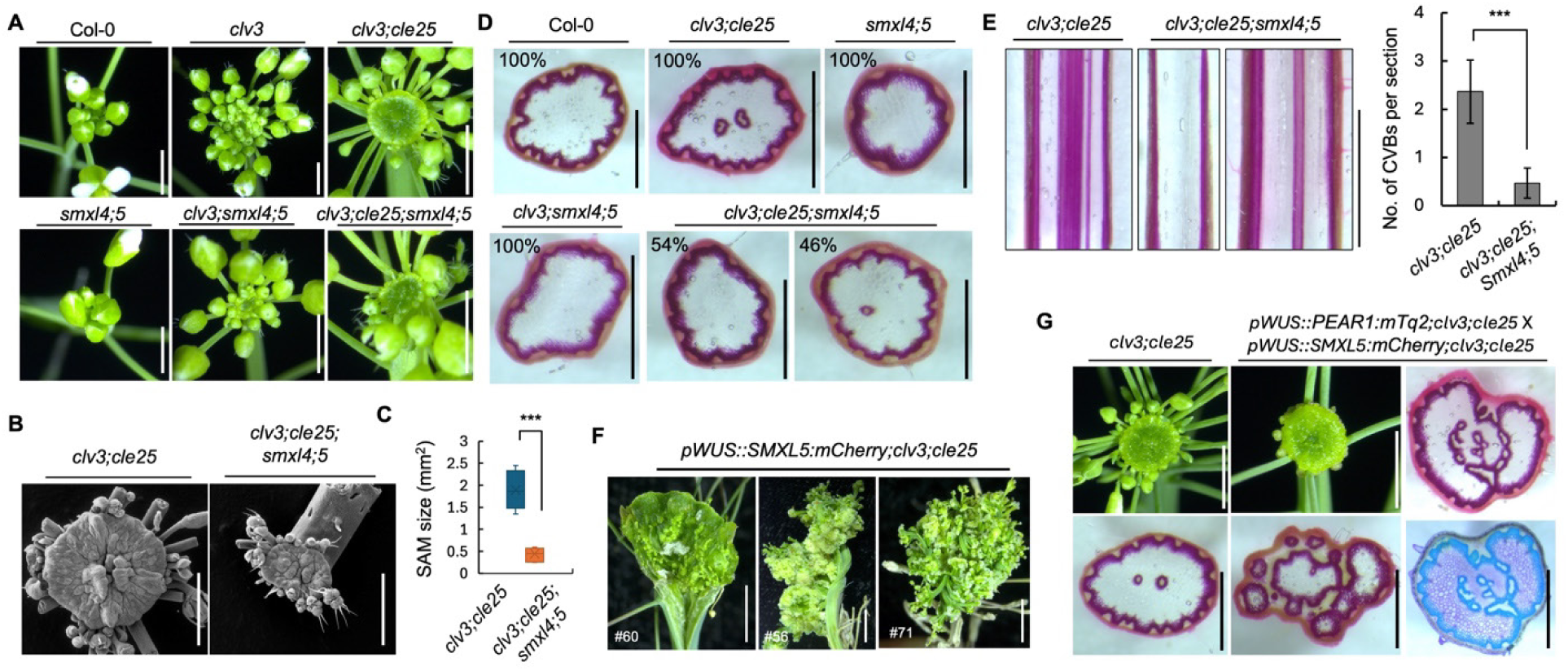
SMXL transcriptional regulators modulate stem cell homeostasis and CVB function in *clv* mutants. (A) Representative top-view SAM images of the indicated genotypes. Scale bars, 2 mm. (B and C) Scanning electron microscopy (SEM) images showing SAM morphology in *clv3;cle25* and *clv3;cle25;smxl4;smxl5* plants (B), and corresponding quantification of SAM diameter (C). Quantification was performed using 6 SAMs for *clv3;cle25* and 7 SAMs for *clv3;cle25;smxl4;smxl5*. Scale bars, 1 mm. Error bars represent SD. Statistical significance: unpaired Student’s t-test, ***P < 0.001. (D) Representative phloroglucinol-stained transverse stem sections of the indicated genotypes. Percentages of plants lacking CVBs and corresponding sample sizes are indicated. Scale bars, 1 mm. (E) Phloroglucinol-stained longitudinal stem sections of mature *clv3;cle25* and *clv3;cle25;smxl4;smxl5* stems (left), and quantification of CVB number (right). Scale bars, 2 mm. Sample size: n = 11 for *clv3;cle25* and n=37 for *clv3;cle25;smxl4;smxl5*. Error bars represent SD. Statistical significance: unpaired Student’s t-test, ***P < 0.001. (F) *pWUS::SMXL5:mCherry;clv3;cle25* transgenic plants showing stem splitting and ectopic accumulation of undifferentiated tissue at shoot apices in mature plants. Scale bars, 1.5 mm. (G) Top-view SAM images highlighting enlarged SAM morphology and altered floral bud number in the indicated F1 plants compared with *clv3;cle25* controls (top left). Corresponding phloroglucinol-stained stem sections reveal disorganization of central and peripheral vascular tissues, and toluidine blue-stained stem sections are shown at right. Scale bars, 1 mm

Importantly, loss of *SMXL4* and *SMXL5* also strongly suppressed CVB development (Figures 6D). Approximately 54% of *clv3;cle25*;*smxl4;smxl5* plants completely lacked detectable CVBs, whereas the remaining plants developed only a single weak vascular structure (Figures 6D and E). Quantitative analysis revealed an average of ∼0.47 CVBs per stem in the quadruple mutant, compared with ∼2.4 CVBs per stem in *clv3;cle25* plants (Figure 6E). These findings indicate that SMXL activity is required for the ectopic CVB formation associated with compromised CLV3p signaling. Together, our results place SMXL proteins downstream of the CLV3p pathway and establish them as critical mediators linking stem cell signaling to vascular bundle patterning.

### SMXLs and PEAR1 cooperatively regulate stem cell maintenance and vascular bundle patterning

Given that PEARs positively regulate *SMXL* transcription (Figure 5A),^29^ we next investigated their coordinated roles in vascular patterning and stem cell maintenance. To this end, we expressed *mCherry*-tagged *SMXL5* under the control of either the *WUS* promoter (*pWUS::SMXL5:mCherry*) or the STM promoter (*pSTM::SMXL5:mCherry*) in Col-0, *clv3*, and *clv3;cle25* backgrounds. Particularly, 2 out of 12 independent *pWUS::SMXL5:mCherry* T1 lines in the *clv3;cle25* background developed ectopic accumulations of undifferentiated tissue at shoot apices during late developmental stages (Figure 6F), phenocopying the abnormalities observed in higher-order CLE peptide–receptor mutants (Figures 3E and S5). These abnormalities were absent in *clv3* backgrounds (Figure S10A). By contrast, *pSTM::SMXL5:mCherry* lines failed to induce comparable stem cell accumulation phenotypes (Figure S11). In addition, neither transgenic population exhibited enhanced CVB formation and instead largely resembled the parental *clv3;cle25* phenotype (Figures S10D and S11), suggesting that overexpression of SMXL5 activity alone has only limited capacity to disturb vascular organization unlike PEAR1 (Figures 5D and S10D). This interpretation is consistent with previous reports showing that SMXL5 activity is regulated post-transcriptionally by *JUL1/2* (*JULGI*)^27^ and relies on additional cofactors, such as OBERON3 (OBE3),^81^ likely due to the absence of an intrinsic DNA-binding domain in SMXL proteins.

To further examine the functional relationship between *PEAR1* and *SMXL5* downstream of CLV3 signaling, we simultaneously overexpressed *PEAR1* and *SMXL5* in the *clv3;cle25* background by crossing the strongest *pWUS::PEAR1:mTq2* and *pWUS::SMXL5:mCherry* transgenic lines. These plants developed severe disk-like SAM structures accompanied by a reduction in floral bud production (Figure 6G). Histochemical analyses revealed extensive vascular abnormalities, including aberrant internal circularization of vascular tissues and disruption of peripheral vascular continuity. Similar defects were observed in toluidine blue–stained stem sections (Figure 6G). Collectively, these results demonstrate that CLV3p signaling maintains vascular bundle organization through coordinated repression of the DOF–SMXL transcriptional module.

### WUS activity directly regulates vascular patterning

Our results indicate that the central *WUS* domain in the SAM serves not only as a core regulator of stem cell maintenance, but also as a key organizer of vascular patterning. Consistent with this model, ectopic activation of phloem-associated DOF transcription factors partially recapitulated *WUS*-related stem cell defects (Figures 5C-E), revealing a functional connection between vascular developmental programs and shoot meristem homeostasis.

To directly investigate the role of WUS activity in VB patterning, we ectopically expressed *WUS* under the *CLV1* promoter in both Col-0 and *clv3* backgrounds. In the *clv3* background, this effectively abolished CLV3p-dependent restriction of *WUS* expression, resulting in expansion of WUS activity beyond its native spatial domain. The *pCLV1::WUS* construct in the Col-0 background did not develop ectopic CVBs. Instead, in the *clv3* background, ectopic *WUS* activity induced the formation of enlarged central pith cells with strong phloroglucinol staining, indicative of elevated lignification (Figure 7A). However, unlike the ectopic CVBs observed in *clv3;cle25*, these structures lacked organized vascular bundle architecture. Toluidine blue staining further revealed that these cells acquired xylem-like characteristics without accompanying phloem differentiation (Figure 7A), indicating that excessive *WUS* activity is sufficient to promote ectopic xylem specification from central pith cells. Collectively, these results establish a direct role for WUS activity in regulating vascular development and stem tissue patterning.

**Figure 7.**
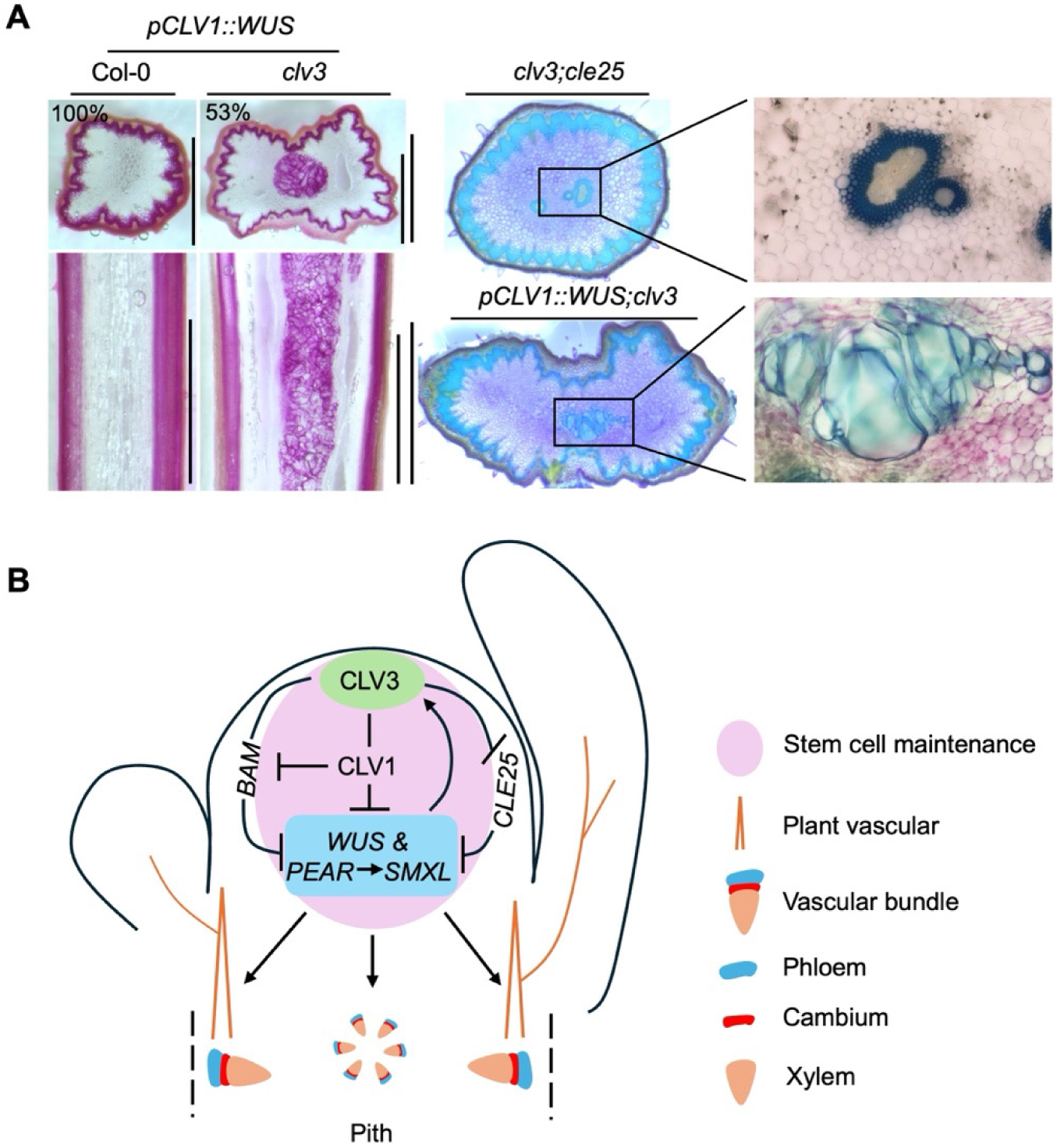
Ectopic WUS activity promotes vascular differentiation in central stem pith cells. (A) Phloroglucinol-stained transverse (top panel) and longitudinal (bottom panel) stem sections of *pCLV1::WUS* transgenic plants in Col-0 and *clv3* backgrounds, highlighting ectopic lignification within the central stem region. Toluidine blue-stained stem sections of *clv3;cle25* and *pCLV1::WUS;clv3* plants are shown on the right. Magnified views of the central stem region are shown. A total of 9 independent T1 plants carrying *pCLV1::WUS* in the Col-0 background and 13 independent T1 plants carrying *pCLV1::WUS* in the *clv3* background were analyzed. Scale bars, 1 mm (transverse sections) and 1.5 mm (longitudinal sections). (B) Schematic model illustrating CLV3p-mediated coordination of stem cell maintenance and vascular development in the stem. Arrows indicate transcriptional activation, T-bars indicate transcriptional repression, and dashed arrows indicate indirect regulatory interactions. Green shading marks the *CLV3* expression domain, light blue shading marks the *WUS* expression domain, and light pink shading denotes the stem cell domain.

## Discussion

How organisms pattern their body is a fundamental question in biology. The correct patterning of stele tissues in plants is essential for plant development and viability. To date most studies on shoot stele patterning have focused on the process of auxin-mediated vascular connection to developing primordia at the stem periphery.^1,3,7,15^ Here we show that the stem cells at the shoot meristem also play a critical role in the patterning of underlying VB, through the production of CLV3p (Figure 7B). CLV3p signaling through overlapping CLV1/BAM receptor suppresses the expression of phloem identity transcriptional network in the SAM center, and consequently the formation CVBs (Figure 1). Notably, this mirrors the role of CLEp signaling in suppressing phloem formation in roots.^58,64,68,74^ Master regulators of phloem formation such as the PEAR and SMXL transcription factor^24,27,29,58^ clades are key targets of CLV3p signaling and are necessary and sufficient to drive ectopic CVBs formation in *clv* mutant shoots (Figure 5D). As such, plants employ a shared peptide based signaling module to shape VB patterning at both poles of the plant body (Figure 7B).

Notably, we find that CLV3p-mediated suppression of CVBs formation also buffers meristem cell proliferation and contributes to classical *clv* stem cell phenotypes (Figures 5 and 6). This occurs via two mechanisms, *CLV3* and *CLV1* transcriptional compensation from paralogues, and the ectopic activity of PEAR and SMXL transcription factors in *clv* SAMs. In Arabidopsis, loss of *CLV1* is transcriptionally compensated by ectopic upregulation of *BAM* genes in the SAM center,^55,67^ while loss of *CLV3* is transcriptionally compensated by ectopic upregulation of *CLE25* (Figure 1G). In both cases, this transcriptional compensation buffers SAM stem cell homeostasis, as higher order mutants with paralogues display enhanced stem cell phenotypes. As neither *clv3* nor *clv1* single mutants display CVB formation (Figures 1F, 3C and D), transcriptional compensation by *CLE* and *BAM* paralogues also masks primary roles for CLV3p-CLV1 signaling in suppression of CVB. Notably, *BAM1* and *BAM2* are expressed in phloem^55^, and *BAM3* and *CLE25* are phloem specific genes.^49,64,68,75^ As such transcriptional compensation in CLV shoot stem cell homeostasis can be explained as a consequence of the partial conversion of the SAM center to phloem tissue in *clv* mutants, resulting in ectopic expression of phloem *CLE* and *BAM* paralogues.

Transcriptional compensation in *clv* signaling in shoot stem cell maintenance is conserved across plant species and was important for crop domestication.^54^ In tomato, loss of *slclv3* is transcriptionally compensated for by *SlCLE9*, whose partial activity in stem cell regulation was permissive for the selection of the beefsteak tomato variety.^54^ Whether this is tied to phloem specification, of if *SlCLE9* is phloem expressed, is unclear. Recently we showed that CLV3p plays a critical role in floral primordia formation in cooler temperatures which is also transcriptionally compensated by CLE25p.^48,56^ In warmer temperatures, heat-induced florigen bypasses the requirement for CLV3p/CLE25p signaling and restores floral primordia generation.^56^ However, we find that warmer temperatures fail to suppress ectopic vascular *in clv3;cle25* mutants, this suggests that CLEp-mediated floral primordia formation and vascular repression are at least partially separable events.

We find that shoot meristem proliferation is also sensitized to the levels *PEAR* and *SMXL* expression in the SAM center (Figures 5 and 6). In wild-type SAMs, CLV3p signaling represses the expression of *PEAR* and *SMXL* transcription factors in the SAM center, which is necessary to prevent ectopic CVB formation (Figure 7B). Increasing the dose of either PEARs or SMXLs in the center of *clv3;cle25* SAMs further enhances the conversion of stem pith tissues into CVB (Figures 5D and 6G). This also enhances stem cell phenotypes in *clv3;cle25* plants, resulting in plants with greatly overproliferated SAM and carpel tissue, fasciation, or increases in carpel numbers, all classical *clv* phenotypes^33,67^ (Figure 5C and E). Correspondingly, mutations in *DOF/PEAR* members suppress carpel numbers in *clv3;cle25*, while *smxl* mutants reduce SAM *clv3;cle25* size (Figures 5C and 6A-C). *dof*/*pear* higher order mutants do not display reduced carpel numbers (Figure 5C), consistent with their repression in SAMs by CLV3p signaling (Figure 1I). As such, phloem transcription factors are partially responsible for classical *clv* meristematic phenotypes, and sufficient to promote *clv*-like proliferation of shoot meristem tissues when expressed ectopically in SAMs (Figures 5E and 6F). Notably, this mirrors the role for PEARs in root stele, where they are necessary and sufficient to drive stele cell proliferation.^29,30,58^ In root stele, phloem DOF/PEAR factors also promote the expression of phloem expressed *CLE* genes (*CLE25*, *CLE26*, *CLE45*) and *BAM3* as part of a negative feedback loop limiting phloem cell proliferation.^58^ In SAMs, ectopic expression of *CLE25* complements *clv3;cle25* mutants (Figure 3A), while ectopic expression of *PEARs* enhances *clv3;cle25* mutant stem cell phenotypes (Figure 5D and E). This implies that PEARs do more in *clv* mutant SAMs than simply promoting paralog transcriptional compensation, again mirroring their role in root stele. It remains to be seen if the targets for PEARs and SMXLs in root and shoot stele are the same, but the shared roles in stele formation, cell proliferation, and the suite of phloem genes repressed by CLV3p suggests a high degree of target overlap across tissues. As in roots, no single factor appears sufficient to promote ectopic stele in wild-type plant SAMs, suggesting this requires the combined activity of multiple factors, with *clv3* and *clv3;cle25* plants being correspondingly sensitized to increases in the levels of stele-promoting factors (Figure 7B).

Strikingly, haplostele-like formation in *clv3* plants is also sensitized to *WUS* dose (Figure 7). While *clv3* mutants do not display ectopic CVB, increasing the expression of *WUS* in *clv3* SAM is sufficient to induce vascular differentiation in stem center (Figure 7A). In this case the stele more closely resembles xylem tissues. This activity mirrors the role of *WUS*-related paralogues *WOX4* and *WOX14*, which promote stele cell proliferation and xylem identity.^43,82,83^ As such, CLV3p signaling from stem cells at the apex restricts the expression of root associated stele networks from the SAM and stem center, with stele transcriptional regulators, or SAM expressed orthologues, playing comparable roles as in root stele. Interestingly, CLV3p mediated repression of CVB formation in the SAM and meristem over-proliferation can be separated through the use of CLV3p variants which discriminate among the different CLEp receptors (Figure 4).^76,77^ This suggests that while diverse CLV1/BAM receptors respond to CLV3p, there may be sub-functionalization of these receptors, as has been suggested during root phloem development.^63,84^

The patterning of stem VB has been a key diagnostic in fossil identification and has been associated with major evolutionary shifts in plant form and function.^10,13,15,22^ Compelling evidence suggests that eustele, with central pith and peripheral VBs, evolved from an ancestral haplostele pattern.^12^ A key question is how pith tissue displaced central VB tissue during evolution. Most evolutionary models consider pith as displacing xylem, but this may be due in part to the comparatively poor preservation of phloem tissues in fossils. The “instrastelar origin hypothesis” posits that central pith arose due to a delay in central VB differentiation, through unknown mechanisms.^6,10,12,14,16,22^ Our data suggests that CLV3p signaling from shoot stem cells could be a plausible mechanism fitting this model (Figure 7B). However, *clv3;cle25* mutants still retain eustele, in addition to a haplostele-like pattern. As such, these two stele tissue types are patterned through partially distinct mechanisms, and derived from separate stem tissues, at least in Arabidopsis. Eustele patterns likely arose from haplostele through a siphonostele intermediates,^12^ which variably display central pith and partially surrounding contiguous stele tissue, disrupted by leaf gaps. It is possible that the evolution of central CLV3p haplostele repression, combined with peripheral auxin mediated stele attraction, could have driven modern eustele evolution. Further exploration will be necessary to assess this central repressor and peripheral attractor model of VB evolution.

## Resource Availability

### Lead Contact

Zachary L. Nimchuk

### Materials Availability

All materials are available upon request to the lead contact.

### Data and Code Availability

Raw transcriptomic data has been deposited into the SRA under project (XXXXXX).

## Acknowledgements

We want to thank Dr. Yrjo Helariutta for providing *pear* mutant and *PEAR* reporter seed lines, Dr. Thomas Greb for providing *smxl* mutant and *SMXL* reporter lines, Dr. Tatsuo Kakimoto for providing *pSUC2::GFP-GUS*, *pCLE25::H2B-GFP:CLE25intron-ter* and *dof-sextuple* lines. The authors want to thank Dr. Pat Gensel for her comments on the manuscript and discussions on the evolution of stelar types. The authors want to thank Jamie Winshell for technical help and expertise, and James Garzoni and John Ward for greenhouse assistance. This work was supported by the U.S. National Institutes of Health (R35GM119614 to Z.L.N) and the U.S. National Science Foundation (IOS-1455607 and IOS-2515446 to Z.L.N.).

## Author contributions

K.R.C. conceptualized and performed experiments, analyzed data, and wrote the manuscript. C.A.S conceptualized and performed experiments and analyzed data. N. P. assisted in plant imaging. Z.L.N. conceptualized the project and experiments, analyzed data, acquired funding, and wrote the manuscript.

## Plant materials and growth conditions

The Arabidopsis thaliana ecotypes Col-0 and Ler were used in this study, with Col-0 serving as the primary genetic background unless otherwise indicated. The *clv3-20*^48^ allele in the Col-0 background was used throughout this study unless otherwise indicated. The following mutant and transgenic lines have been described previously: *clv3-2* (*Ler* background), *cle25,*^48^ *clv3;cle25,*^48^ *cle16;cle17,*^71^ *clv3-15;cle16,*^71^ *clv3-15;cle*17,^71^ *cle33;cle45,*^74^ *clv3-15;cle16;cle17,*^71^ *clv3;cle26*,^48^ *clv3;cle27*,^48^ *clv3;cle26;cle27*,^48^ *clv3;cle25;cle26*,^48^ *clv3;cle25;cle27*,^48^ *clv1-15,*^48^ *clv1-4*^38^ (*Ler* background), *bam1,*^67^ *bam3,*^67^ *bam1;bam2,*^67^ *bam1;bam2;bam3,*^67^ *cik1;cik2;cik4,*^48^ *cik1;cik3;cik4;cik5,*^48^ *cik2;cik3;cik4,*^48^ *cik1;cik2;cik3;cik4;cik5;cik6,*^48^ *dof-sextuple* (*obp2;dof2.2;pear1;pear2;dof5.3;dof3.*2),^58^ *smxl4-1,*^24^ *smxl5-1,*^24^ *smxl4;smxl5,*^24^ and the reporter lines *pCLE25::H2B-GFP:CLE25intron-ter,*^58^ *pSUC2::GFP-GUS,*^58^ *pPEAR1::VENUSer,*^29^ *pPEAR2::VENUSer,*^29^ *pPEAR2::PEAR2:VENUS,*^29^ *pOBP2::YFP,*^29^ *pSMXL4::YFP,*^24^ and *pSMXL5::YFP*.^24^

Seeds were surface sterilized in 70% ethanol containing 0.05% SDS for 10 min, washed three times with 100% ethanol, and plated on half-strength Murashige and Skoog (MS) medium supplemented with appropriate selection antibiotics where appropriate.^85^ Seeds were stratified in darkness at 4°C and subsequently transferred to long-day growth conditions (16 h light/8 h dark) at 22°C. Nine-day-old seedlings were transplanted to soil (propagation mix supplemented with perlite and Peter’s 20:20:20 [N:P:K] fertilizer at recommended concentrations).

### DEX inducible *CLV3* construct design, induction, and RNA-Seq

A previously described *DEX::CLV3* construct^37^ was introduced into the *clv3-2* and *clv1-4;clv3-2* backgrounds to generate *DEX::CLV3;clv3-2* and *DEX::CLV3;clv1-4;clv3-2* transgenic lines, respectively. As a control, a *DEX::mTq2* construct was introduced into the *clv3-2* background. Plants were grown under standard growth conditions, and when the inflorescence stems reached 1 to 4 cm in height, plants were sprayed with either mock solution or 10 μM dexamethasone (DEX). For *DEX::CLV3;clv3-2* and *DEX::CLV3;clv1-4;clv3-2* lines, shoot apical meristems (SAMs) were harvested 5 h or 24 h following treatment. Floral buds were removed immediately prior to collection, and SAMs were flash-frozen in liquid nitrogen. For *DEX::mTq2;clv3-2* control plants, SAMs were harvested 24 h after mock or DEX treatment. Samples were stored at −80°C until RNA extraction. Three independent biological replicates were collected for each condition, with each replicate consisting of 100 pooled inflorescence meristems.

Total RNA was extracted using the RNeasy Plant Micro Kit (Qiagen; cat no 74034) according to the manufacturer’s instructions. RNA-seq libraries were prepared and sequenced on the BGIseq-500 platform. Sequencing reads were aligned to the *Arabidopsis thaliana* reference genome (TAIR10) using STAR, and transcript abundance was quantified using Salmon. Differential gene expression analysis was performed using standard RNA-seq workflows as described earlier.^48^

### CRISPR constructs and mutagenesis

To generate *CLV3* and *CLE25* mutations in the *dof-sextuple*^58^ background, CRISPR–Cas9-mediated genome editing was performed using 19- and 20-bp gene-specific single-guide RNAs (sgRNAs), respectively, as previously described for the generation of *clv3;cle25* lines.^48^ The zCas9i cloning system, which contains an intron-optimized GFP-tagged Cas9, was used for all constructs.^86^ Previously published guide sequences targeting *CLV3* (ATGAAAATGGAAAGTGAAT) and *CLE25* (GGTTGGAGTGATTGCATCTT) were used.^48^ The respective sgRNAs were first cloned into the shuttle vectors *pDGE332* and *pDGE334* using a modified Golden Gate assembly with *BpiI* and subsequently assembled into the destination vector *pDGE667* using *BsaI-HFv2.*^87^ Final constructs were sequence verified and introduced into the *dof-sextuple* background via Agrobacterium-mediated transformation using the floral dip method. T1 transformants were selected on hygromycin-containing medium and genotyped for Cas9 by PCR. Cas9-free T2 plants were selected based on seed coat FAST-Red fluorescence and further confirmed by PCR genotyping. Mutations in *CLV3* and *CLE25* were validated by Sanger sequencing and subsequently used for phenotypic and molecular analyses.

### Scanning electron microscopy (SEM)

For SEM analysis, young inflorescences were dissected and floral buds were carefully removed without damaging the shoot apical meristem (SAM). Samples were fixed in 100% methanol, which was gradually exchanged with 100% ethanol prior to critical point drying (CPD). CPD was performed according to the manufacturer’s instructions. Dried samples were mounted onto specimen holders using double-sided conductive carbon adhesive tabs (25 mm OD PELCO Tabs; Ted Pella). Imaging was performed using a Hitachi TM4000Plus II scanning electron microscope equipped with a motorized stage and operated in charge-reduction vacuum mode.

### Plasmid constructs and transgenic lines

To generate the *pCLV3::CLV3*, *pCLV3::CLE25*, *pSUC2::CLE25*, and *pATML1::CLV3* constructs, coding sequences (CDSs) of *CLV3* and *CLE25* were TOPO-cloned into the *pENTR-D* vector (Thermo Fisher Scientific, K243520) as previously described in our laboratory.^48^ For the *pCLV3::CLE45* and *pATML1::CLE45* constructs, the full-length *CLE45* CDS was synthesized and cloned into the pTwist-ENTR vector by Twist Biosciences (https://www.twistbioscience.com). Entry clones were subsequently recombined into binary destination vectors by LR Clonase-mediated Gateway cloning (Thermo Fisher Scientific). The *CLV3* promoter vector (*MOA33pCLV3*) was used to generate *pCLV3::CLV3*, *pCLV3::CLE25*, and *pCLV3::CLE45*; the *SUC2* promoter vector (*MOA34pSUC2*)^56^ was used to generate *pSUC2::CLE25*; and the *ATML1* promoter vector (*MOA33pATML1*) was used to generate *pATML1::CLV3* and *pATML1::CLE45*. As a control construct, *YPet* was expressed under the *ATML1* promoter to generate *pATML1::YPet*.

To generate *WUS* promoter-driven constructs, CDSs of *SMXL4*, *SMXL5*, and *PEAR1* fused to *mCherry* or *mTq2* fluorescent tags were synthesized and cloned into *pTwist-ENTR* vectors by Twist Bioscience (https://www.twistbioscience.com) to generate the entry constructs *pTwist-ENTR SMXL4:mCherry*, *pTwist-ENTR SMXL5:mCherry*, and *pTwist-ENTR PEAR1:mTq2*. Entry clones were subsequently recombined into binary destination vectors by LR Clonase-mediated Gateway cloning. The *MOA33pWUS* destination vector was used to generate *pWUS::SMXL5:mCherry*, whereas the *MOA34pWUS* destination vector was used to generate *pWUS::SMXL4:mCherry* and *pWUS::PEAR1:mTq2*. The *SMXL5:mCherry* entry construct was additionally recombined into the *STM* promoter vector (*MOA34pSTM*) to generate *pSTM::SMXL5:mCherry*. Similarly, the full-length *DOF2.2* CDS was fused to *mTq2* and cloned into *pTwist-ENTR* and recombined into *MOA34pSTM* to generate *pSTM::DOF2.2:mTq2*. To generate *pCLV1::WUS*, the *WUS* CDS was TOPO-cloned into *pENTR-D* and subsequently recombined into the *CLV1* promoter destination vector (*MOA34pCLV1*).

*MOA33*-based vectors confer kanamycin resistance, whereas *MOA34*-based vectors confer hygromycin resistance.^88^ All binary constructs were transformed into *Agrobacterium tumefaciens* strain GV3101 by electroporation and introduced into plants by the floral dip method. Transgenic lines were generated in the indicated genetic backgrounds, and T1 transformants were selected on the appropriate antibiotics and further validated by PCR genotyping. Primer sequences are listed in Data S1. T1 and T2 generation plants were used for imaging and downstream analyses.

### Stem sections and histochemical staining

Fully mature first and second internodes from the main stems of the indicated genotypes were used for histochemical analysis of vascular tissues. Stems were sectioned using a razor blade to obtain thin transverse and longitudinal sections, which were then stained with either phloroglucinol–HCl or toluidine blue. Phloroglucinol–HCl (Wiesner) staining solution was prepared by dissolving 0.3 g of phloroglucinol in 10 mL of absolute ethanol to obtain a 3% (w/v) stock solution.^89^ The solution was mixed thoroughly, and one volume of 37 N HCL was added to two volumes of the phloroglucinol solution, followed by additional mixing, as described earlier.^89^ Freshly prepared phloroglucinol–HCl solution was used for all analyses. Toluidine blue O staining solution (0.02%) was prepared by dissolving 0.02 g toluidine blue O in 100 mL distilled water and stored for repeated use.^89^ Stem sections were transferred to 2.0 mL microcentrifuge tubes and incubated for 1–2 min in the respective staining solutions. Stained sections were imaged using a Zeiss Stemi 2000-C equipped with a Zeiss Axiocam 105 color or Nikon Eclipse 80i compound microscope using DIC optics.

### GUS staining

GUS reporter activity was analyzed following established procedures.^90^ Stems were thinly hand-sectioned using a laser blade and immediately collected in ice-cold 90% acetone. Samples were then rinsed with GUS staining buffer containing 50 mM sodium phosphate (pH 7.0), 0.2% Triton X-100, 5 mM potassium ferrocyanide, and 5 mM potassium ferricyanide to reduce background staining and limit diffusion of reaction products. After washing, tissues were incubated in freshly prepared staining buffer supplemented with 2 mM X-Gluc and maintained at 37°C until the appearance of detectable blue coloration.^90^ Enzymatic activity was quenched by exchanging the staining solution with 70% ethanol. Samples were subsequently imaged using a Zeiss Stemi 2000-C equipped with a Zeiss Axiocam 105 color.

### Peptide treatment and root growth assay

CLV3, CLV3^P4R^, and CLE45 peptides were synthesized by Biomatik (http://www.biomatik.com) and dissolved in water according to the manufacturer’s instructions. Each peptide was applied at a final concentration of 0.1 µM on ½-strength MS agar plates, with a no-peptide treatment used as a control.^62^ Seedlings of the indicated genotypes were grown on the respective media, and root lengths were imaged and quantified.

### Site Directed Mutagenesis of CLV3

Wild-type *CLV3* CDS cloned into the *pENTR-D* entry vector (Thermo Fisher Scientific, K243520), as previously described in our laboratory, was used as the template for site-directed mutagenesis. Point mutations corresponding to the *CLV3^P4R^* and *CLV3^P4R/P7S^*variants were introduced using the Q5® Site-Directed Mutagenesis Kit according to the manufacturer’s instructions (NEB E0554S). The resulting mutant entry clones, *pENTR-D CLV3^P4R^* and *pENTR-D CLV3^P4R/P7S^*, were confirmed by Sanger sequencing. Sequence-verified mutant CDSs were subsequently recombined into the binary destination vector *MOA33pCLV3* by LR Clonase-mediated Gateway cloning to generate the final expression constructs *MOA33pCLV3::CLV3^P4R^* and *MOA33pCLV3::CLV3^P4R/P7S^*. Primer sequences are listed in key resources table.

### RNA isolation and RT-qPCR

Young inflorescence meristems from the indicated genotypes were harvested when the shoot apical meristem (SAM) contained approximately 7–12 floral buds. Floral buds were removed, and ∼50–70 mg of SAM-enriched tissue was collected per biological replicate (n = 3). Tissue was homogenized using a mortar and pestle, and total RNA was extracted using RNAzol RT with 4-bromoanisole and GlycoBlue, according to the manufacturer’s instructions. First-strand cDNA was synthesized from 2 µg of total RNA using the ProtoScript II First Strand cDNA Synthesis Kit. Quantitative RT-PCR (qRT-PCR) was performed using PowerUp SYBR Green Master Mix on an Applied Biosystems QuantStudio 6 Flex system. Relative transcript levels were calculated using the 2^−ΔΔCT method, with *PP2A* serving as the internal reference gene.^91^ Primer sequences are listed in key resources table.

### Microscopy imaging

For confocal microscopy, shoot apical meristems (SAMs) from long-day–grown plants of the indicated genotypes were harvested when stems carried approximately 5–10 floral buds. Floral buds were first removed in cold water following embedding of the stem in 2% agarose at the bottom of 35 mm plastic dishes, as previously established in our laboratory. The dissected SAMs were then incubated in 1.5 mM propidium iodide (PI) prepared in PIPES buffer (35 mM PIPES, 2 mM MgSO₄, 5 mM EGTA, pH 6.8) for 1–5 min depending on genotype, followed by two washes in fresh water.^48,56^ Petri dishes with prepared SAMs were then filled with water and imaged on a Zeiss LSM980 upright confocal microscope with a 20x/1.0 NA water-dipping lens. Confocal stacks were converted into maximum-intensity projections using ImageJ for figure preparation. Acquisition and image processing settings were kept constant across all SAMs for a given reporter line, whereas PI signal intensity was adjusted per sample to optimize structural visualization.

## Supplemental Figures and Figure Legends

**Figure S1.**
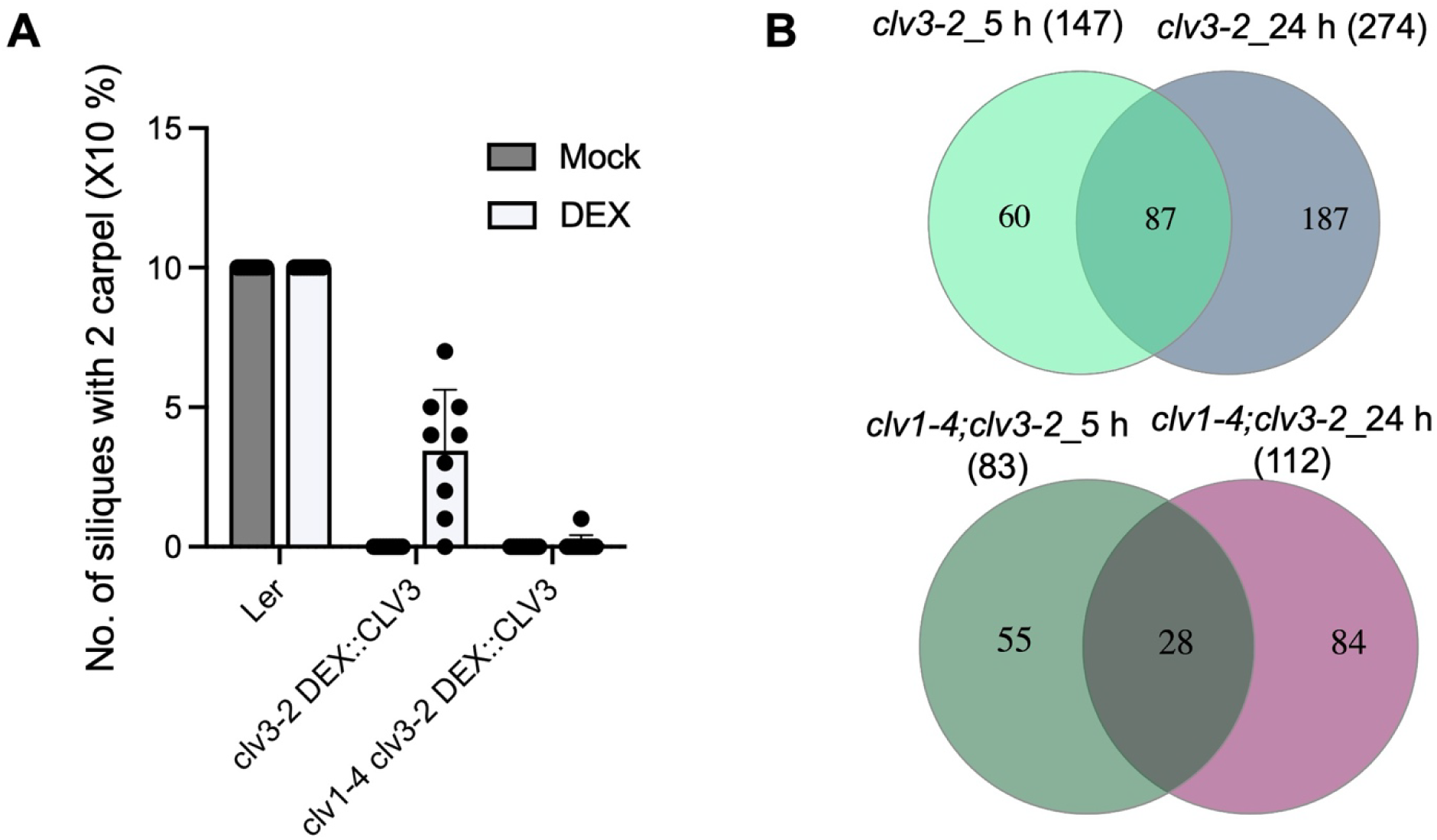
Carpel quantification and transcriptomic analysis following DEX-inducible *CLV3* activation. (A) Quantification of carpel number, represented as the proportion of siliques containing two carpels, upon mock or 10 ìM dexamethasone (DEX) treatment in the indicated genotypes. *clv3-2* and *clv1-4* are in the Ler background. For each genotype, 8–10 siliques from 7–10 independent plants were scored, and mean values are shown. Error bars represent SD. (B) Venn diagrams showing overlap among differentially expressed genes identified by RNA-seq following DEX-induced *CLV3* activation in the *clv3-2;DEX::CLV3* (top panel) and the *clv1-4;clv3-2* mutant background (bottom panel) after 5 h and 24 h of treatment. Numbers indicate unique and shared differentially expressed genes between conditions.

**Figure S2.**
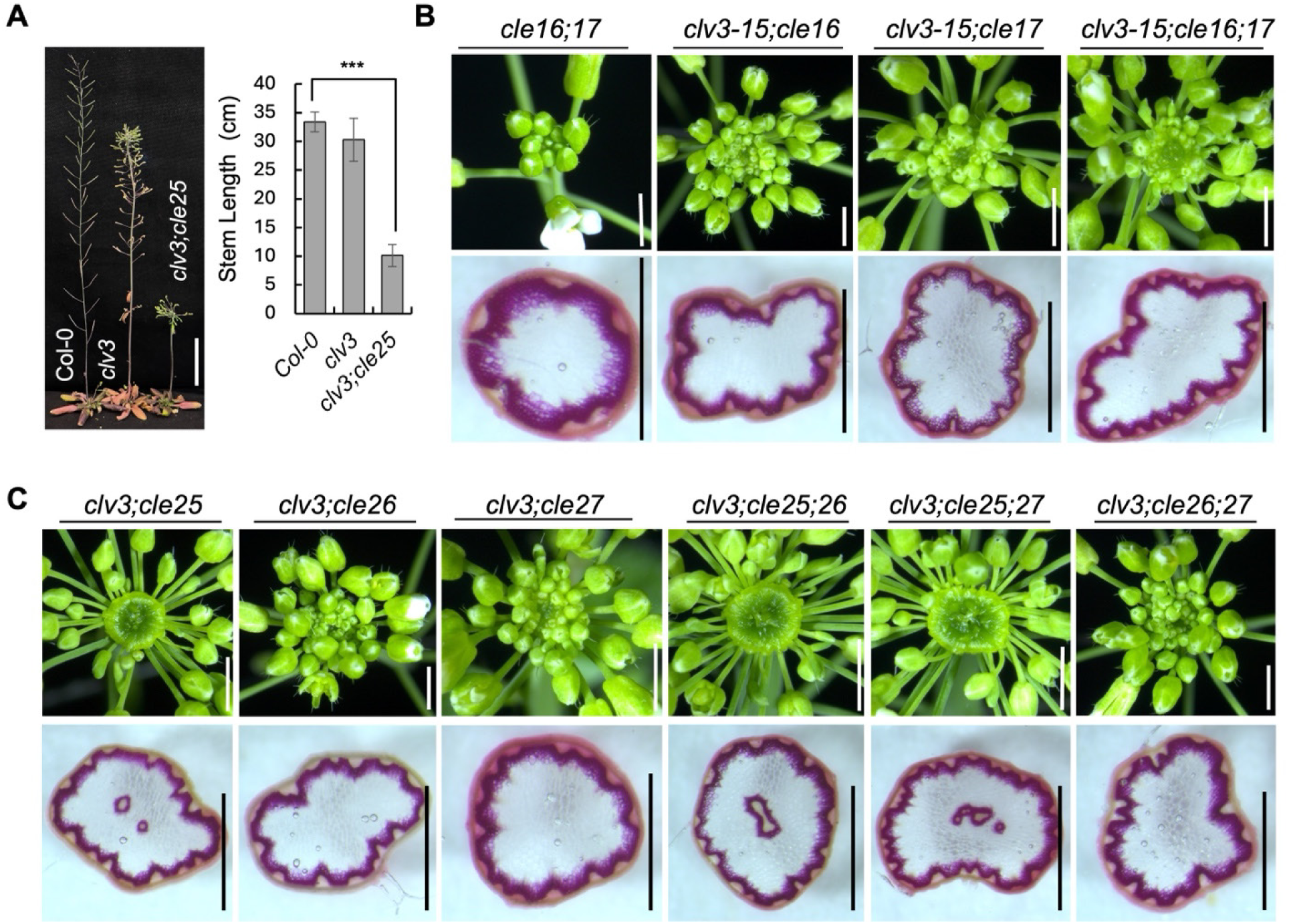
CLV3p and CLE25p function in the suppression of CVB formation. (A) Two-month-old plants (left) and mean stem length of the indicated genotypes. Scale bar, 5 cm. Sample size, n = 7–10; error bars, SD. Statistical significance was determined using an unpaired Student’s t-test (***P < 0.001). (B) SAM images (top panel) of the indicated genotypes and corresponding phloroglucinol-stained stem sections (bottom panel). Scale bars, 2 mm for SAM images and 1 mm for stem sections. (C) SAM images (top panel) and corresponding phloroglucinol-stained stem sections (bottom panel) showing SAM disc-like morphology and central vascular bundle (CVB) formation in the indicated *cle* mutant combinations. Scale bars, 2 mm for SAM images and 1 mm for stem sections.

**Figure S3.**
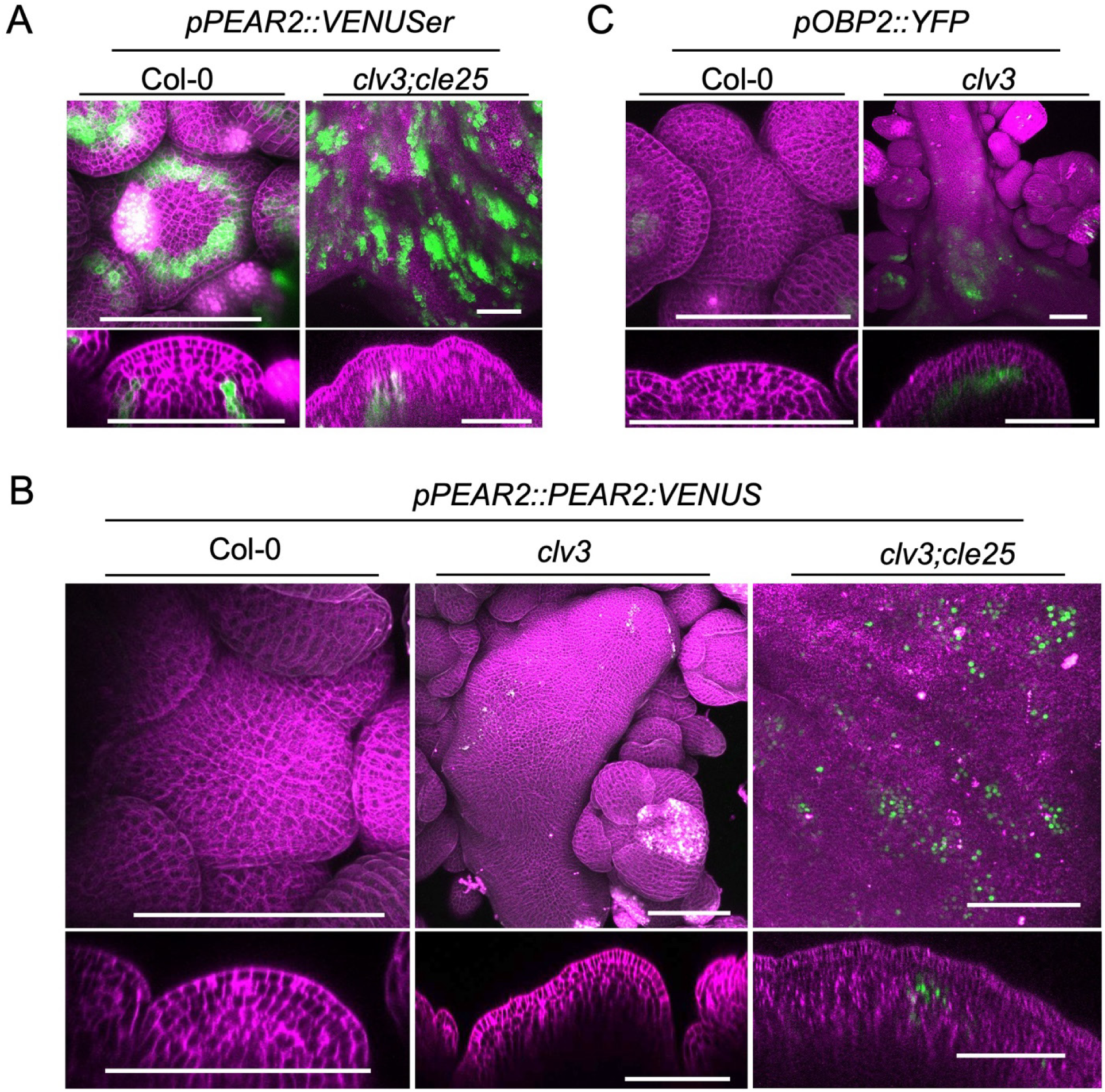
Expression analysis of phloem-DOF transcription factor reporters. **(A** and **B)** Maximum-intensity projections of *pPEAR2::VENUSer* (A), and *pPEAR2::PEAR2:VENUS* (B) expression in the indicated shoot apical meristems (SAMs). Vertical views (top panels) and longitudinal optical sections (bottom panels) are shown. Scale bars, 100 μm. Representative images from three biologically independent samples are shown. (B) Maximum-intensity projections of *pOBP2::YFP* expression in Col-0, and *clv3* SAMs. Vertical views (top panels) and longitudinal optical sections (bottom panels) are shown. Scale bar, 100 μm. Representative images from three biologically independent samples are shown.

**Figure S4.**
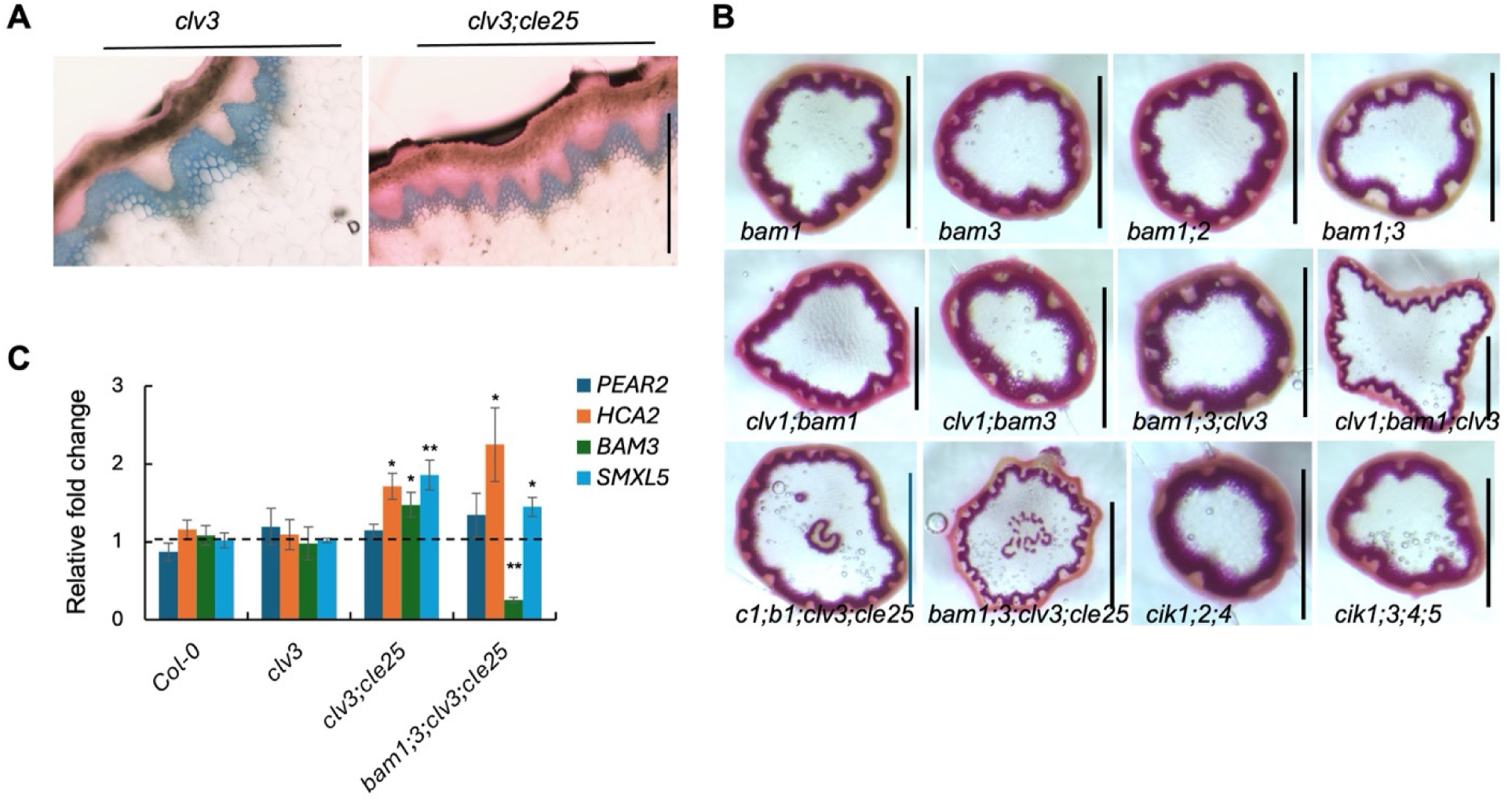
CLE receptors are required for suppression of CVB formation. (A) Toluidine blue–stained magnified stem images of *clv3* and *clv3;cle25*, highlighting increased peripheral vascular bundles in *clv3;cle25*. Scale bar, 0.1 mm. (B) Phloroglucinol-stained stem sections of CLE receptor mutant combinations. *c1;b1;clv3;cle25* is indicated as *clv1;bam1;clv3;cle25*. (C) Relative phloem gene expression measured by RT–qPCR in SAM tissues of the indicated genotypes. *PP2A* was used as the internal control. Error bars represent SD; dotted line indicates control (Col-0) levels. Sample size, n = 3. Statistical significance was determined using an unpaired Student’s t-test. Asterisks indicate statistical significance, with P < 0.05 (*) and P < 0.01 (**)

**Figure S5.**
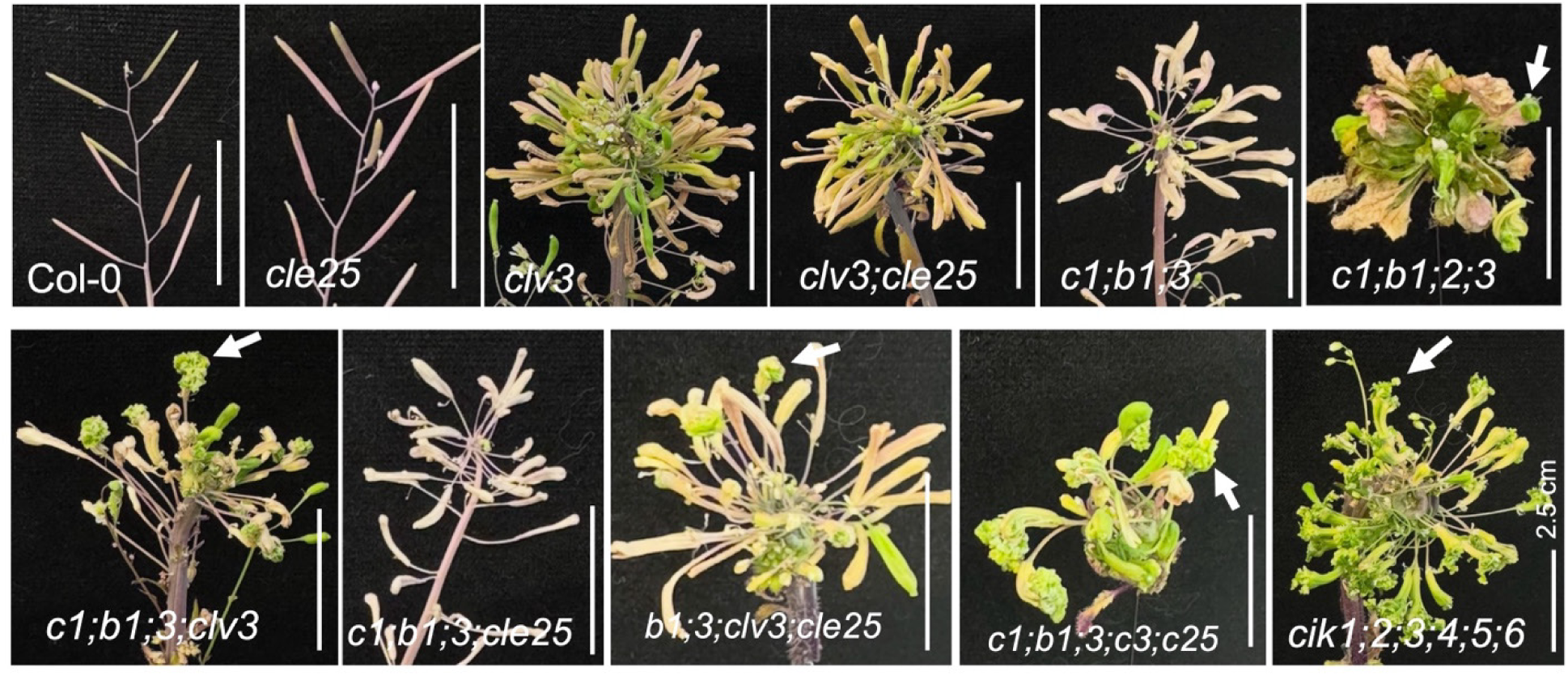
Floral tips from two-month-old plants showing accumulation of undifferentiated tissue at carpel tips (arrows). *c1* indicates *clv1*, *b1* indicates *bam1*, *b2* indicates *bam2*, *b3* indicates *bam3*, *c3* indicates *clv3*, and *c25* indicates *cle25*. Scale bar, 2.5 cm.

**Figure S6.**
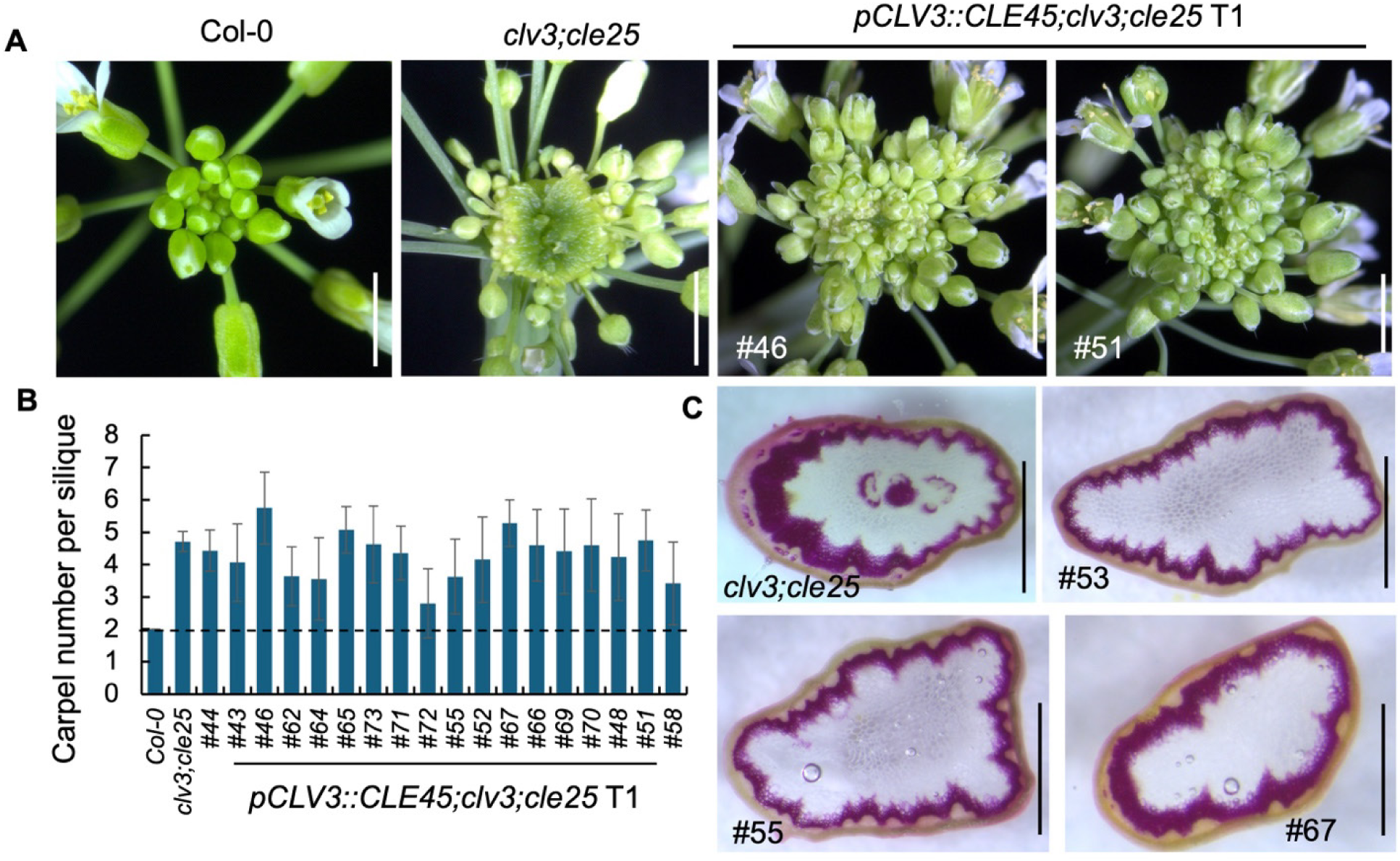
Ectopic expression of CLE45 decouples suppression of central vascular bundle (CVB) formation from stem cell phenotypes. (A) SAM images of the indicated genotypes showing the disc-like phenotype of *clv3;cle25* and its partial suppression in two independent T1 lines of *pCLV3::CLE45;clv3;cle25*. Scale bars, 2 mm. (B) Quantification of carpel number in Col-0, *clv3;cle25*, and 18 independent T1 lines of *pCLV3::CLE45;clv3;cle25*. Error bars represent SD; dotted line indicates the Col-0 carpel number. (C) Phloroglucinol-stained stem sections of *clv3;cle25* and three independent T1 lines (#53, #55, and #67) of *pCLV3::CLE45;clv3;cle25*. Scale bars, 1 mm.

**Figure S7.**
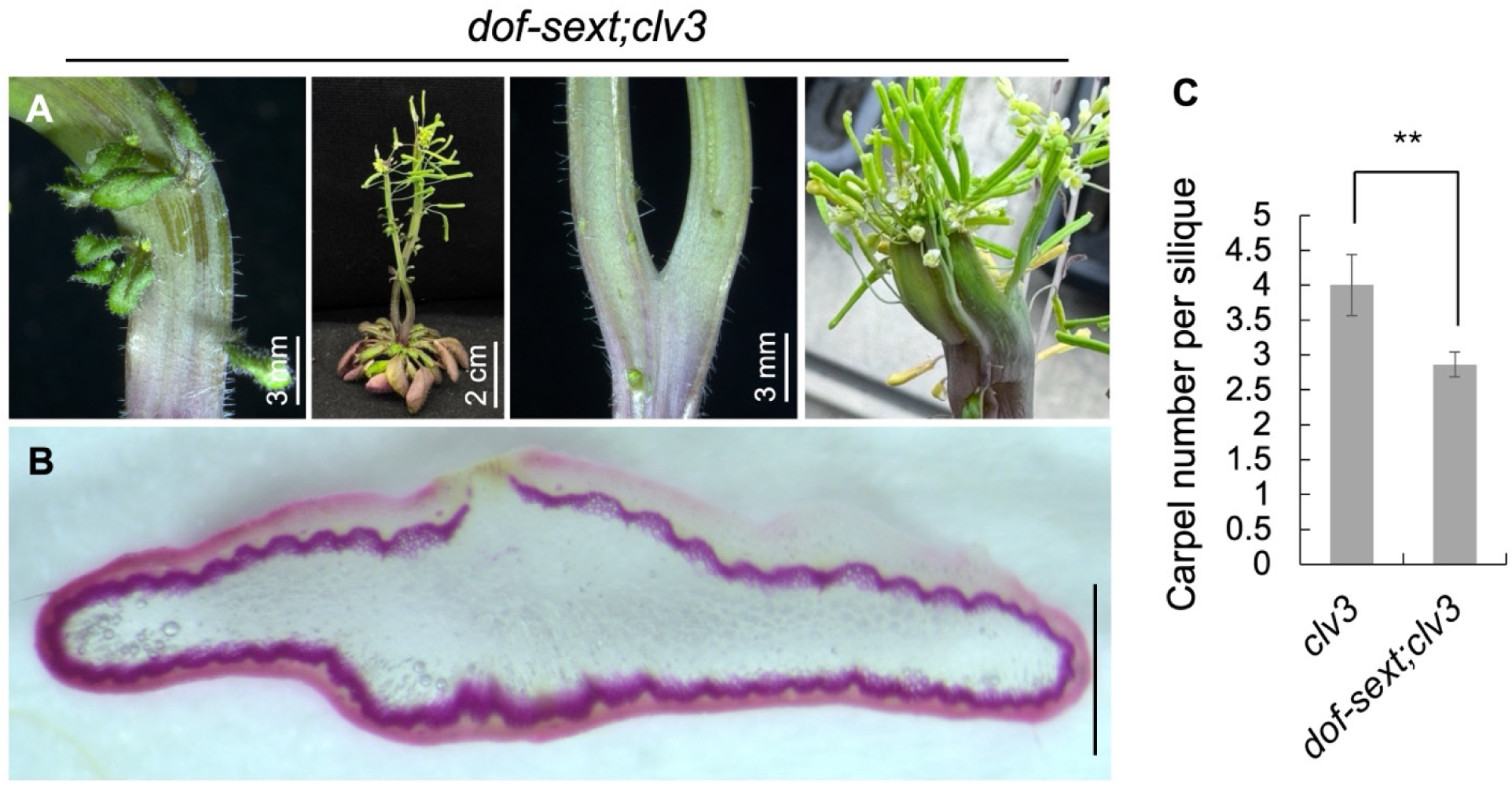
Phenotypic analysis of *dof-sext;clv3*. (A) Two-month-old plants of *dof-sext;clv3* showing widened and bifurcated stem phenotypes. Scale bars are indicated. (B) Phloroglucinol-stained stem sections of *dof-sext;clv3*. Scale bar, 1 mm. (C) Quantification of mean carpel number per silique in the indicated genotypes. For each genotype, 10–15 siliques were analyzed from 4 independent plants. Error bars represent SD. Statistical significance was assessed using an unpaired Student’s *t*-test; ** indicates *P* = 0.008.

**Figure S8.**
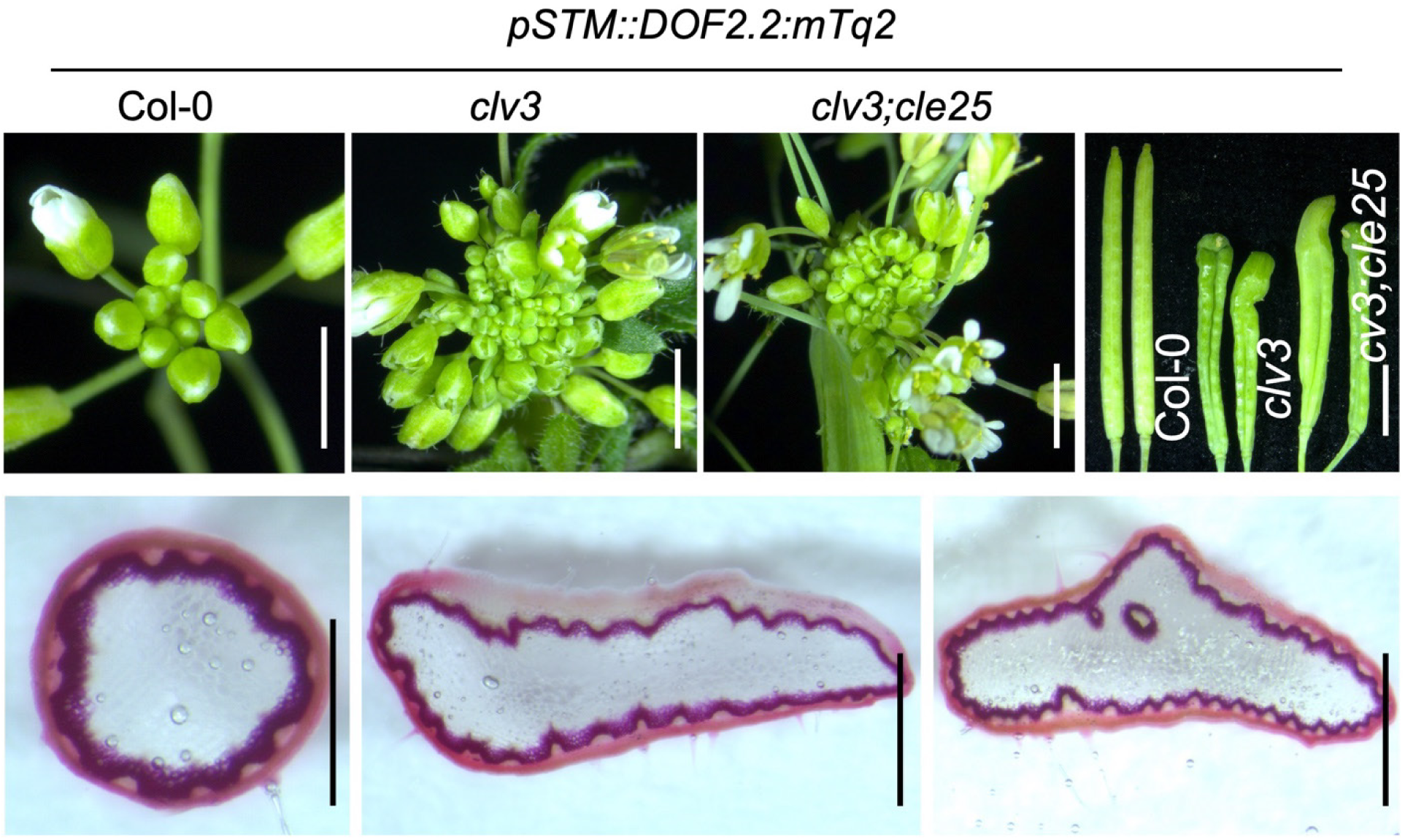
Ectopic expression of *DOF2.2* under the *STM* promoter. Top-view SAM images (top panel) and corresponding phloroglucinol-stained stem sections (bottom panel) of T1 plants expressing *pSTM::DOF2.2:mTq2* in Col-0, *clv3*, and *clv3;cle25* backgrounds. Carpel images of the indicated transgenic lines are shown in the top-right panel. Scale bars, 2 mm (top panel) and 1 mm (bottom panel). Sample size, n = 6 independent T1 plants per genotype.

**Figure S9.**
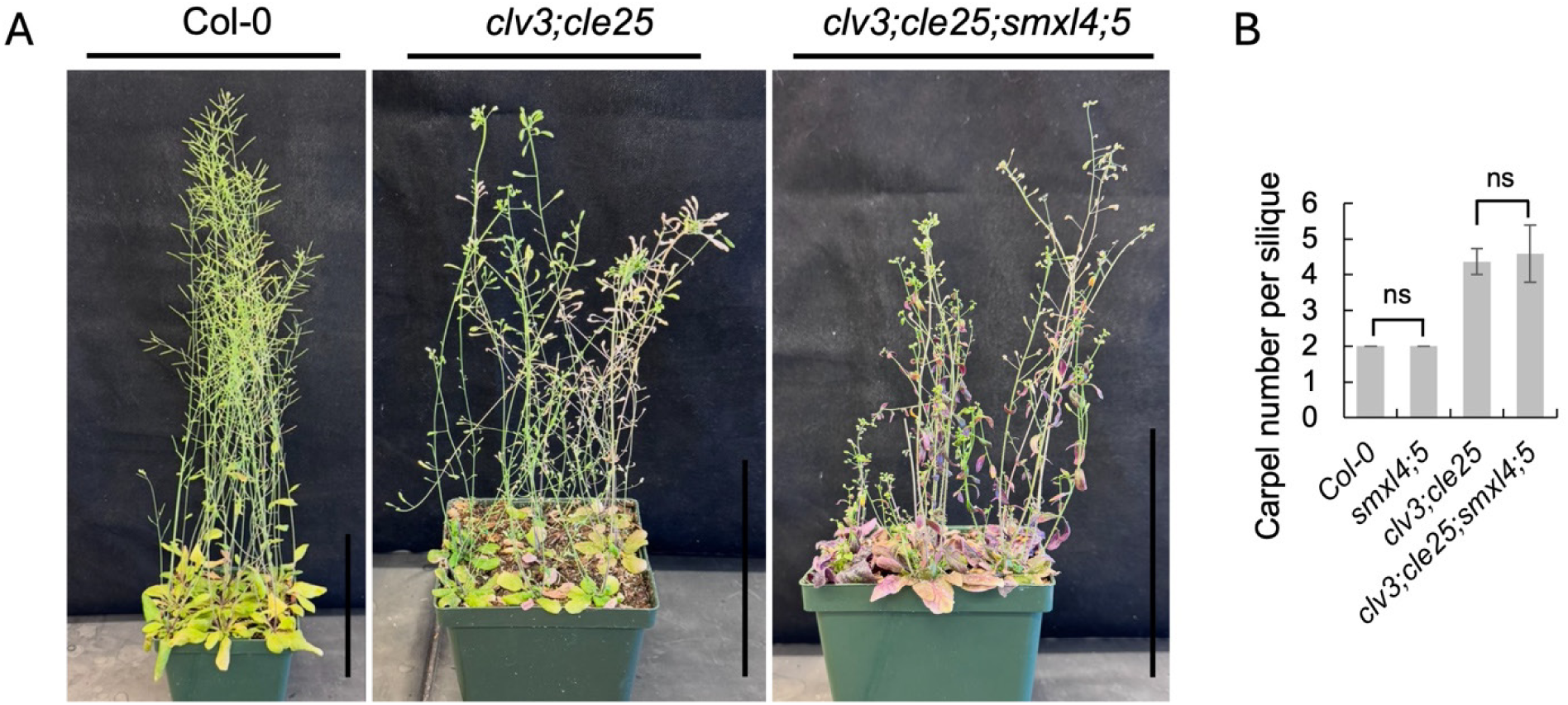
Genetic interaction between *clv3;cle25* and *smxl4;smxl5*. (A) Mature plants of the indicated genotypes. *clv3;cle25;smxl4;smxl5* plants display enhanced anthocyanin accumulation, reduced fertility, and decreased stem height. Scale bars, 10 cm. (B) Quantification of carpel number in the indicated genotypes. For each genotype, 8–10 siliques per plant were scored from 5–7 independent plants. *smxl4;5* indicate *smxl4;smxl5*. Error bars represent SD. Statistical significance was determined using an unpaired Student’s t-test; ns, not significant.

**Figure S10.**
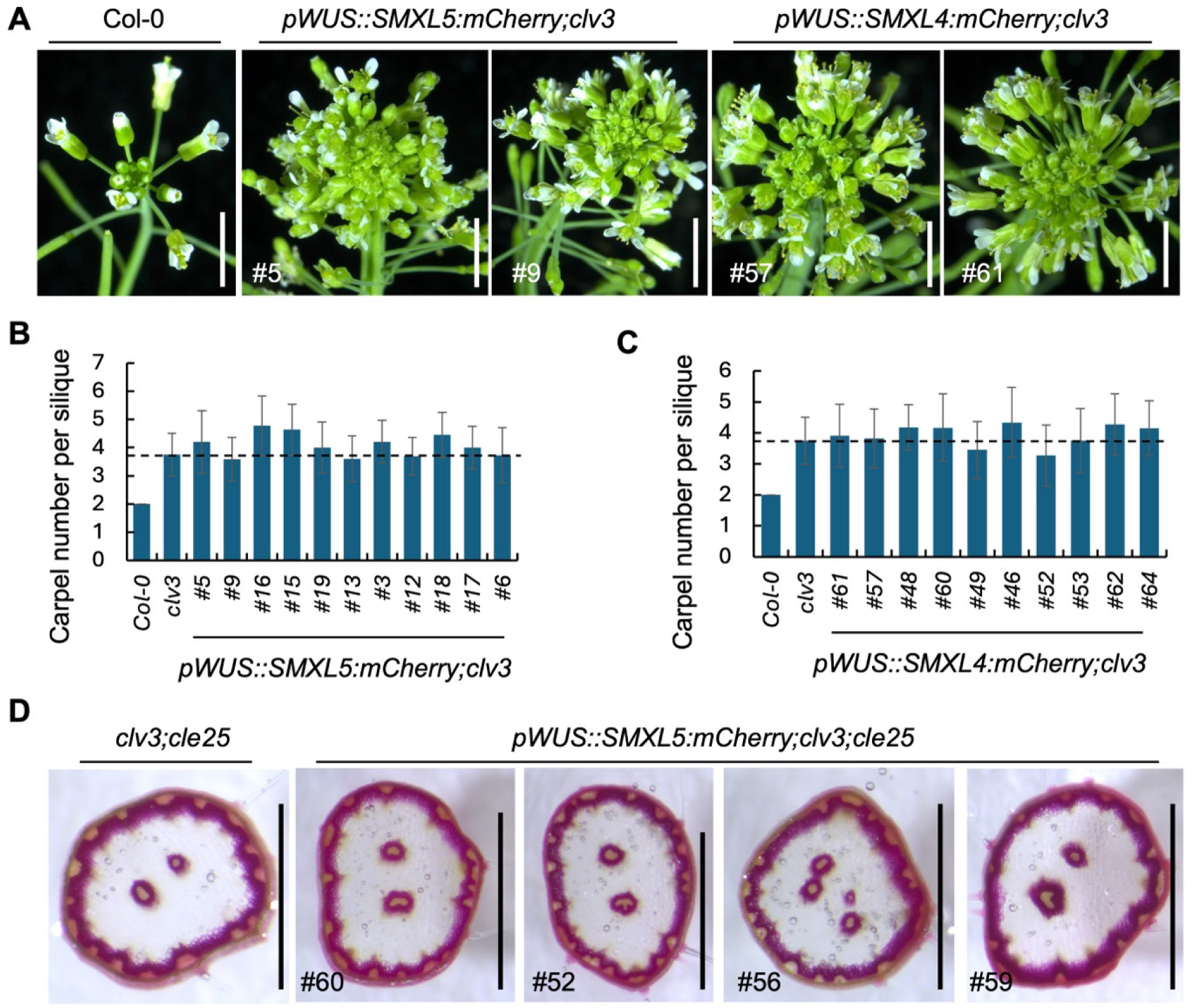
Ectopic expression analysis of *SMXL4* and *SMXL5*. (A) SAM images of Col-0 and T1 plants of the indicated genotypes. Corresponding line numbers are shown. Note: *pWUS::SMXL4:mCherry*; and *pWUS::SMXL5:mCherry* construct transformed into the *pCLE25::H2B-GFP:CLE25intron-ter;clv3* background. Scale bars, 2 mm. (B and C) Quantification of average carpel number in the indicated T1 lines together with Col-0 and *clv3* controls. Note that the same Col-0 and *clv3* carpel number datasets were used in both (B) and (C), as plants were grown and analyzed simultaneously. Note: *pWUS::SMXL4:mCherry*; and *pWUS::SMXL5:mCherry* construct transformed into the *pCLE25::H2B-GFP:CLE25intron-ter;clv3* background. Error bars represent SD. Dotted lines indicate the corresponding *clv3* carpel number. A total of 8–12 carpels were analyzed for each genotype. (D) Phloroglucinol-stained stem sections showing vascular organization in the indicated T1 plants. Corresponding line numbers are shown. Scale bar, 1 mm.

**Figure S11.**
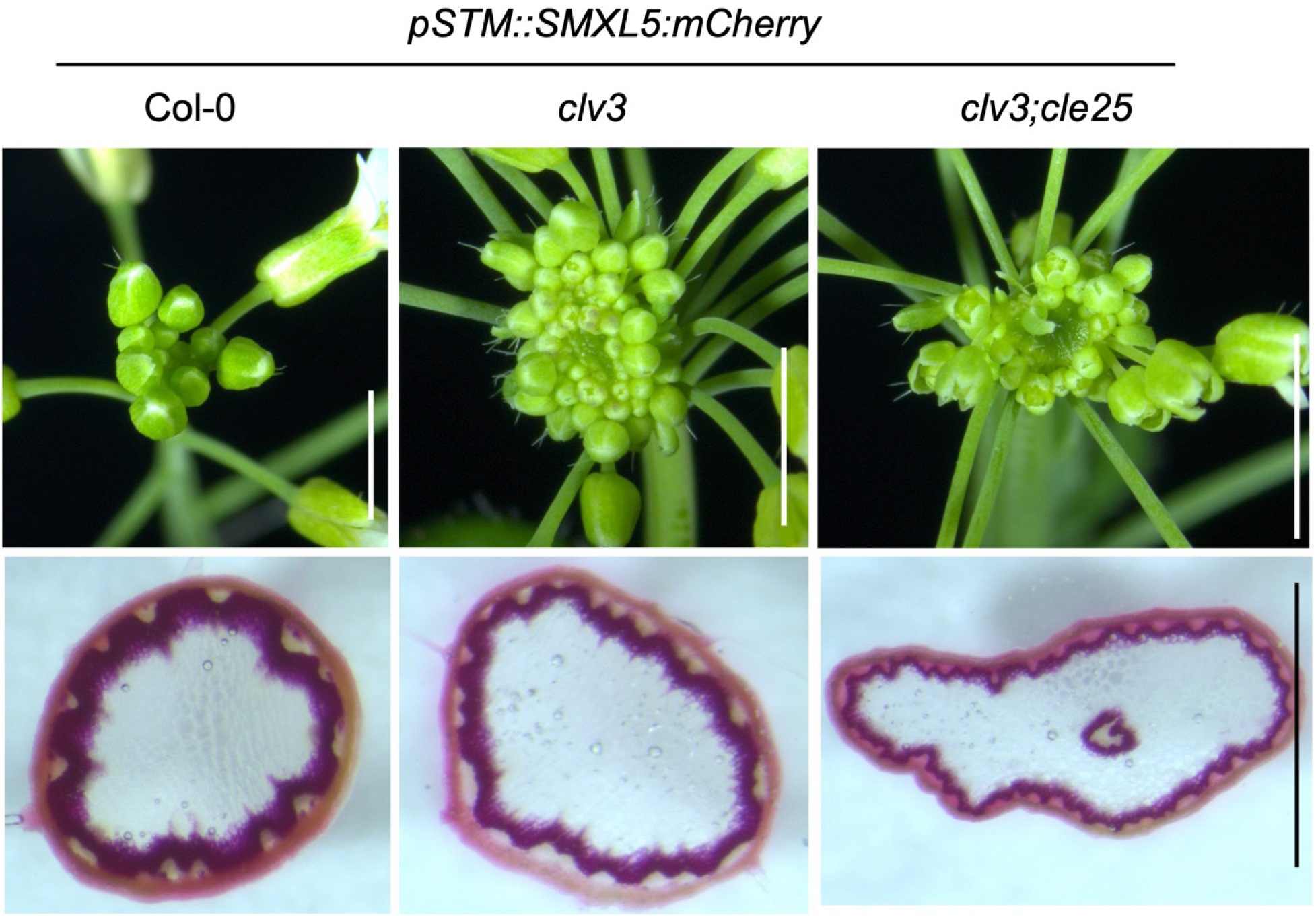
Functional analysis of *SMXL5* misexpression in the *STM* domain. SAM images (top panel) and corresponding phloroglucinol-stained stem sections (bottom panel) of T1 plants expressing *pSTM::SMXL5:mCherry* in Col-0, *clv3*, and *clv3;cle25* backgrounds. Scale bars, 2 mm (top panel) and 1 mm (bottom panel). Sample sizes: *pSTM::SMXL5:mCherry* Col-0 (n = 6), *pSTM::SMXL5:mCherry;clv3* (n = 4), and *pSTM::SMXL5:mCherry*;*clv3;cle25* (n = 7).

## Key Resources Table

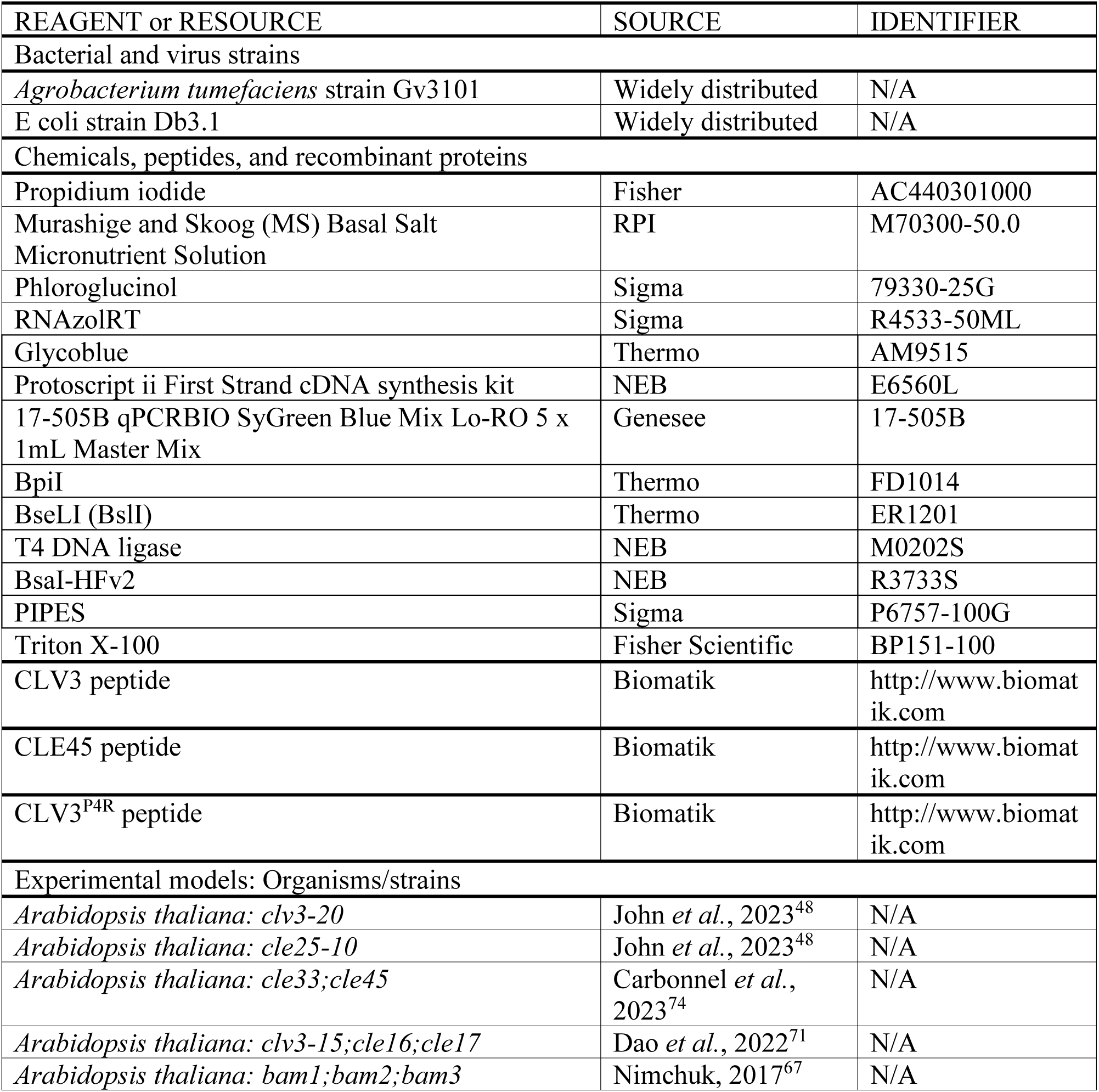
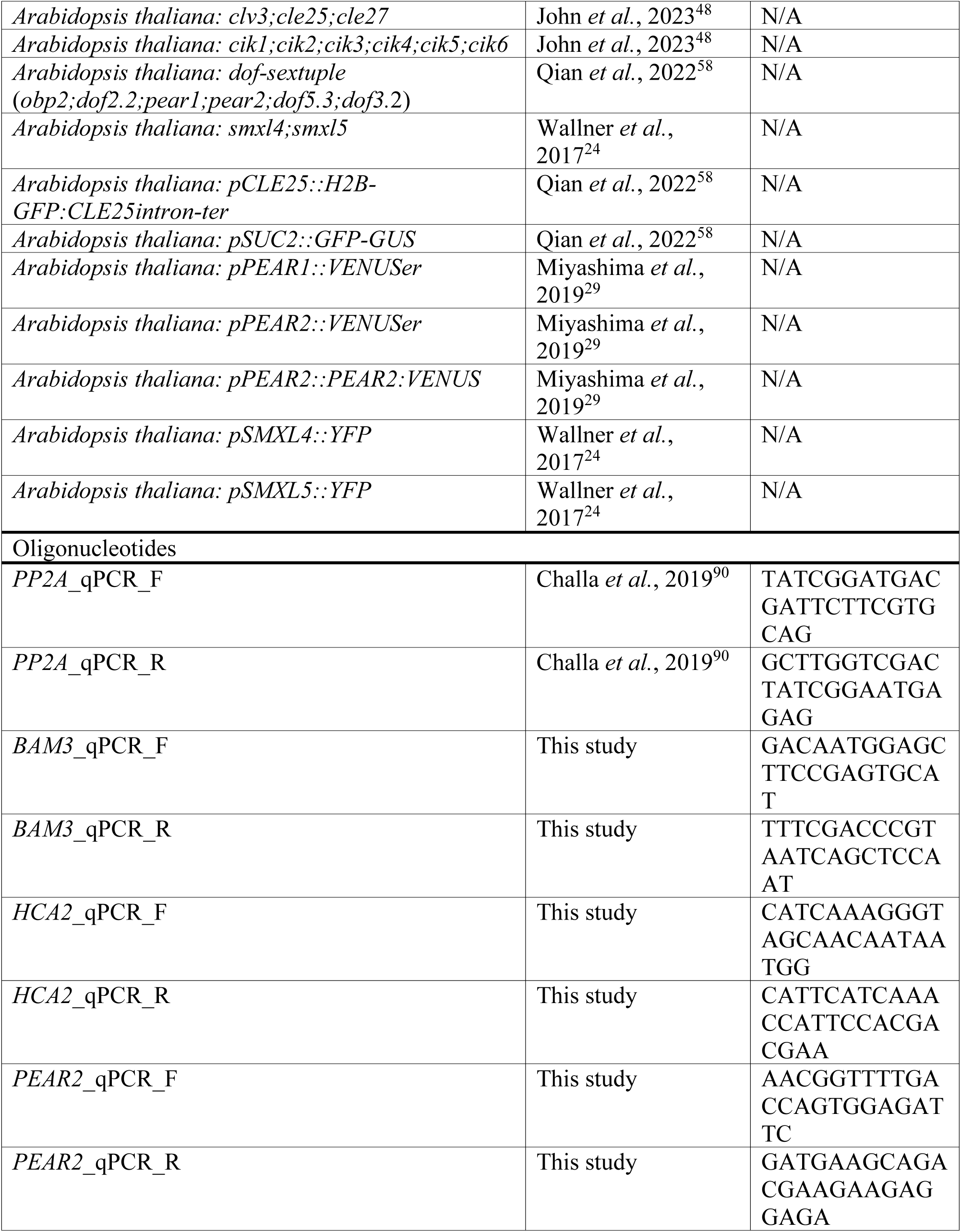
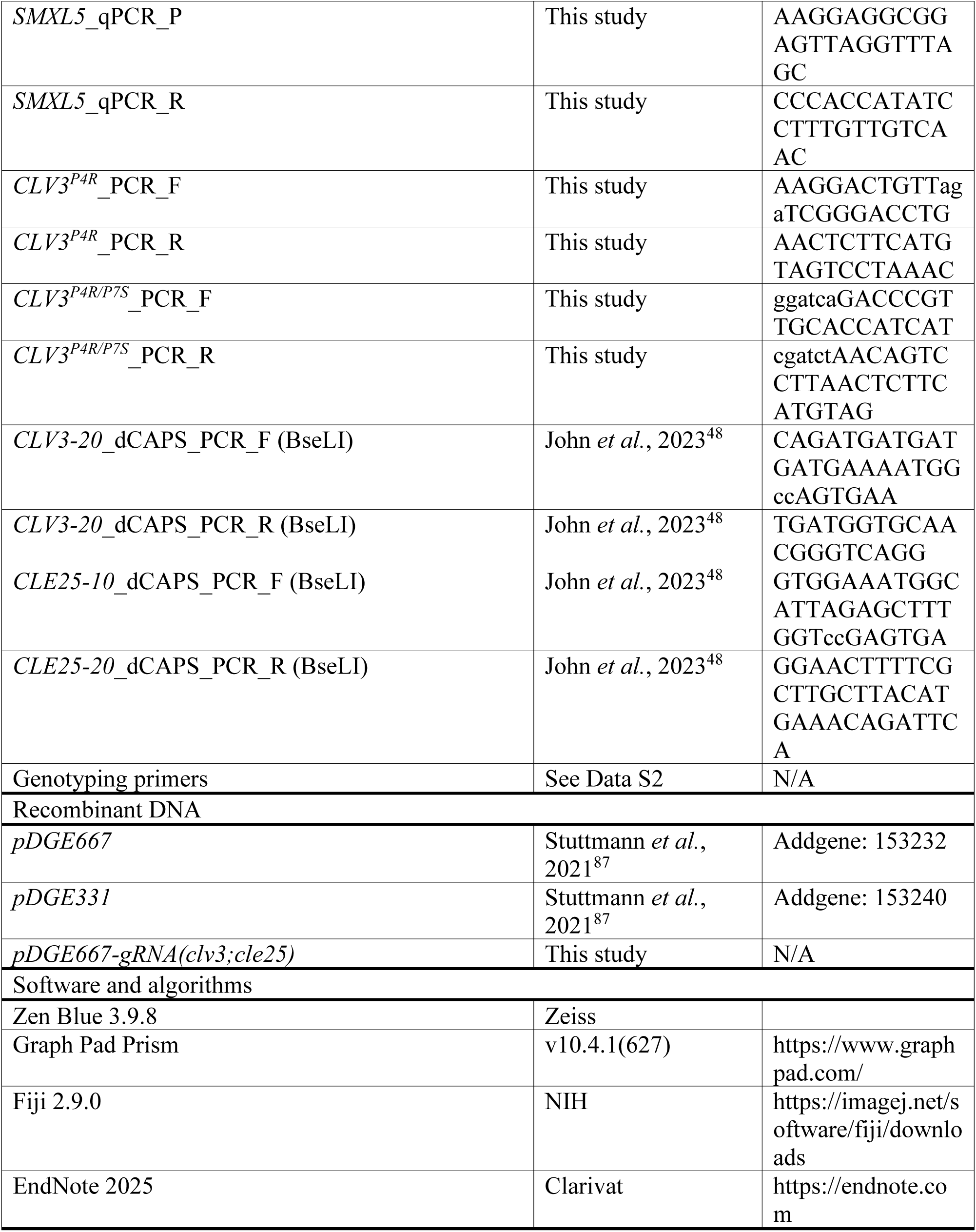

